# Hierarchically engineered multi-enzyme nanoreactors for *in vitro* drug biosynthesis and pathway transplantation into cells

**DOI:** 10.64898/2026.02.15.704820

**Authors:** Ainur Sharip, Somayah S. Qutub, Manar M. Farooqui, Walaa Baslyman, Nida Khalfay, Lukman O. Alimi, Patricia Lopez Sanchez, Lingyun Zhao, Milena Chernyshevskaia, Giovanni Colombo, Niveen M. Khashab, Stefan T. Arold, Raik Grünberg

**Author notes:** contributed equally (Ainur Sharip, Somayah S. Qutub).

## Abstract

Even though most proteins evolved to function within multi-protein systems such as metabolic networks or signaling pathways, most technical protein applications are based on isolated proteins, for example therapeutic antibodies or industrial enzymes. Reliable methods for the stabilization, *in vitro* operation and intracellular delivery of multi-protein systems could unlock new applications in green biotechnology, diagnostics, and medical therapy. We demonstrate that the entire violacein biosynthesis pathway, consisting of up to six separately purified enzymes, can be infiltrated into a hierarchically etched MIL-101 (eMIL) metal organic framework. eMIL nanoreactors reshaped pathway dynamics and reaction flows, multiplied violacein yield *in vitro* and enabled pathway reuse, lyophilization and storage. Moreover, eMIL nanoreactors delivered the six-protein system into mammalian cells, where it produced violacein from cell-provided substrates and cofactors. These findings pave the way for the design of “smart” stimuli-responsive multi-enzyme nanoreactors for biotechnological and medical applications.

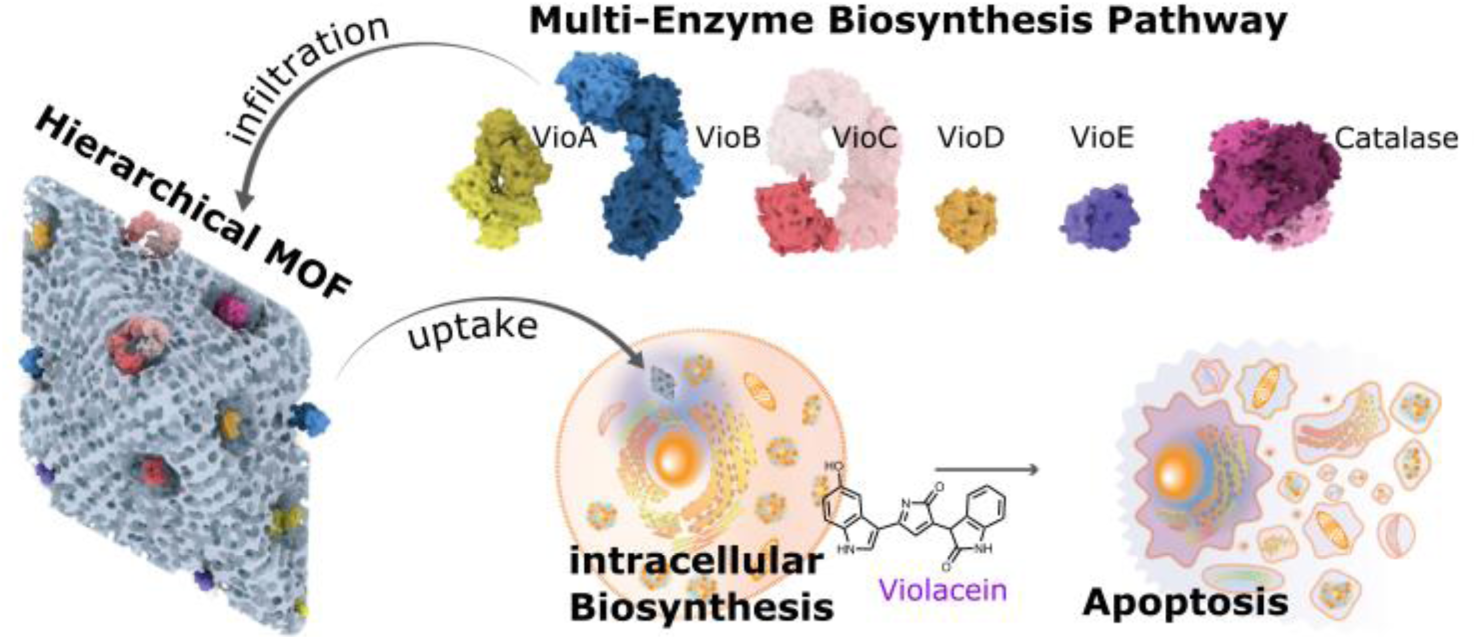

## 1. Introduction

A $300 billion market testifies to the large impact of protein therapies. Current applications are dominated by recombinant antibodies and hormone therapies while therapeutic enzymes^1^ are playing a steadily increasing role. However, protein therapies are held back by at least three important limitations:

Firstly, established protein drugs can only target receptors on the cell surface or in extracellular spaces. Methods for the effective delivery of proteins across the cell membrane and into the cell are therefore actively sought after. The fusion to cell-penetrating peptides, the packaging into micelles, vesicles or protein cages or the attachment to various kinds of nanoparticles have shown promise^2^ but, in practice, the cell interior remains the domain of more traditional small molecule drugs. A second limitation of protein therapeutics is their inherent low stability. Whereas traditional drugs are conveniently handled and stored at ambient temperatures, the typical protein is formulated in liquid and depends on an uninterrupted cold chain during its limited shelf life. Such requirements drive up costs and often exclude remote or less privileged populations^3^. A third limitation is of both technical and conceptual nature: Biologicals in use today are based on a single active protein (such as one specific antibody or hormone). By contrast, most proteins have evolved to perform their function in concert with others. For instance, most small molecule drugs derive from natural products^4^ which originate from multi-step biosynthesis pathways embedded within larger metabolic networks. Such multi-enzyme systems achieve more complex chemistry and can sense and respond to a cell’s metabolic needs. In contrast to a small molecule drug or single protein agent, the intracellular delivery of multi-protein systems could potentially intervene in biological networks with higher fidelity and some degree of dynamic control. Nevertheless, there appears to be no precedent for the delivery of a functionally synergistic multi-protein system into any type of cell. So far, at most three proteins have been co-delivered with a coated zeolitic imidazolate frameworks-8 (ZIF-8) metal-organic framework (MOF); yet these proteins were non-cooperating proof-of-concept molecules^5^.

MOFs had previously emerged as promising materials for *in vitro* enzyme immobilization^6,7^. MOFs are hybrid crystalline porous materials composed of metal ion nodes connected by organic linker molecules^8^. The tunability of the size and structure of MOF’s pores can create versatile matrices for the entrapment of enzymes. The resulting nano- to micrometer-sized enzyme-loaded particles have a very large surface area and high diffusibility of substrates and products. Proteins can be entrapped by diffusion into the pores of assembled MOFs^9–11^, however, in most cases, this requires secondary enlargement of micropores to facilitate entry of the large enzymes. Another approach is the self-assembly of the MOF around the protein, which, however, is only possible for those MOFs that crystallize under conditions that do not denature proteins^12–16^. MOF encapsulation often enhances a protein’s stability under harsh conditions without precluding its catalytic activity^17^. The co-encapsulation of up to three model enzymes (β-galactosidase, glucose oxidase, horseradish peroxidase) has been demonstrated in ZIF-8^18^ but native multi-enzyme pathways have not yet been incorporated into MOFs, even for *in vitro* use.

Some MOFs can deliver their cargo directly into cells. Initial studies demonstrated the delivery of small molecules^19,20^, in particular anti-cancer drugs^21^. However, MOFs are now also used for the intracellular delivery of functional proteins^22–31^. Examples include the delivery and endosomal release of a CRISPR Cas9 endonuclease complex^22^, the aforementioned delivery of three model proteins^5^, delivery of antigens for vaccination purposes^23^, or the delivery of DNA polymerases for miRNA detection^25^ or DNA computation^31^. In contrast to more established protein transfection methods (such as microinjection, liposomes or electroporation) that only apply *ex vivo*^32,33^, MOF delivery translates to animal models where biodegradable MOFs are often well tolerated^34^.

Herein, we report the successful incorporation of the complete biosynthesis pathway for the natural product violacein, relying on up to six enzymes, into a modified MOF Material Institute Lavoisier-101 (MIL-101) (**Figure 1**). MOF infiltration increases product yield and renders the pathway reusable and more robust for *in vitro* synthesis reactions. Moreover, we demonstrate the delivery of the entire multi-protein system into mammalian cells, where it interacts with the specific metabolic state of cancer cells leading to the enhanced production of the cytotoxic violacein as an *in-situ* therapeutic.

**Figure 1.**
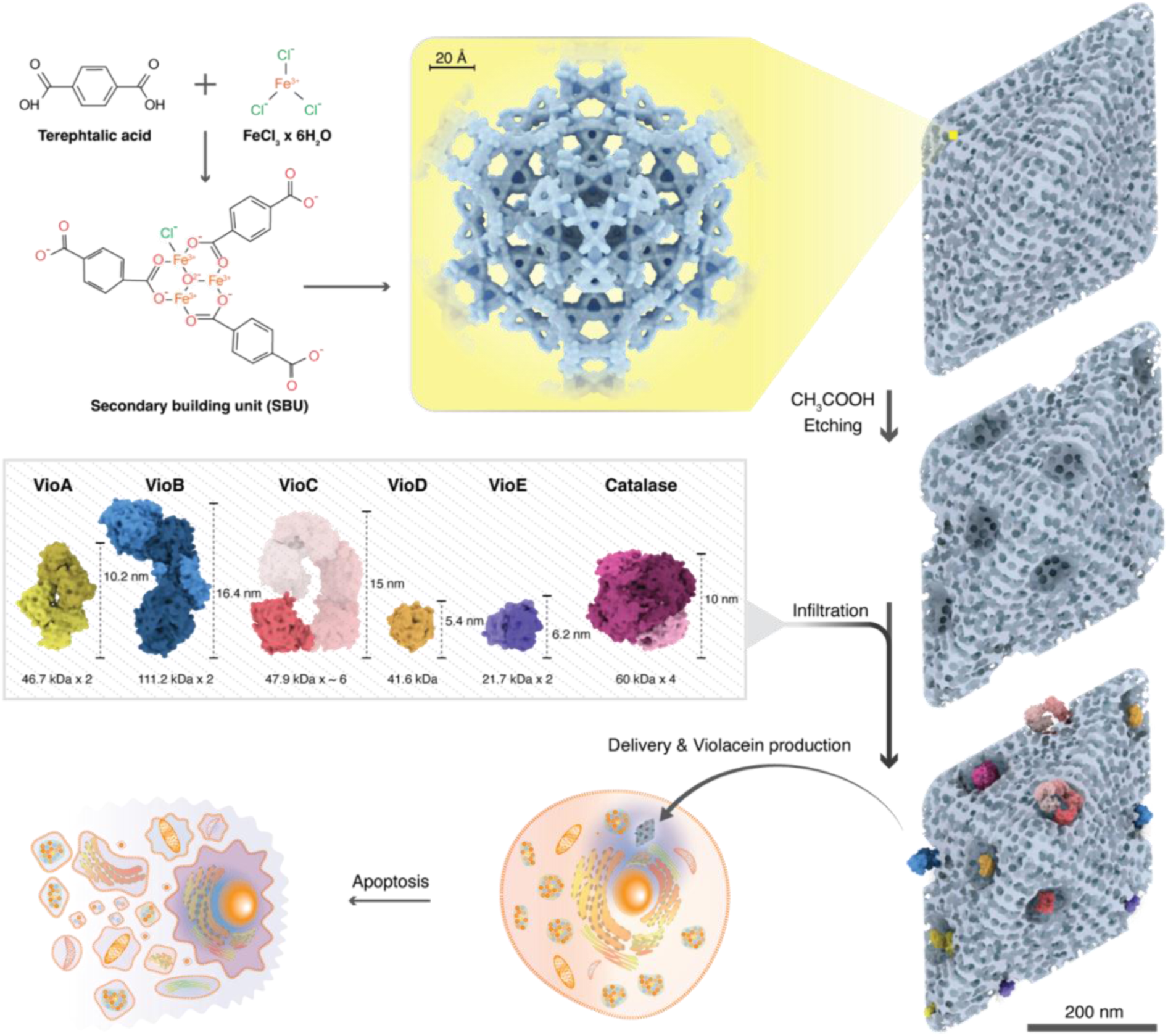
Construction of multi-enzyme nanoreactors. The pore size of unmodified MIL-101^35^ (1.5 nm, yellow box) is too small for the infiltration of the violacein pathway enzymes which have sizes between 5 and 16 nm and some of which adopt oligomerization states (gray box, further detail in Figure 3 and the text). The etching of larger holes allowed for the co-infiltration of individually purified proteins. Pathway-loaded MOF nanoreactors produce violacein *in vitro* but can also be delivered into mammalian cells for *in situ* production of the natural product, leading to apoptosis.

## 2. Results

### 2.1. *In vitro* reconstitution of the violacein pathway

Violacein is a natural purple-colored product that was first isolated from *Chromobacterium violaceum*, although other violacein-producing microorganisms have since been identified^36,37^. Violacein production is controlled by the quorum sensing system to counter competing bacteria, fungi, nematodes or viruses^38^. Violacein has attracted considerable interest owing to its medical and pharmacological activities with demonstrated antimicrobial, antiparasitic, antiviral, and antiprotozoal effects. In addition, it was shown to have immunomodulatory, analgesic, antipyretic, anti-diarrheal and ulcer-protective functions^39^. The violacein pathway is encoded within an operon of five genes (*vioABCDE*). These genes translate into five enzymes (VioA to VioE) that together catalyze the formation of violacein from two molecules of tryptophan and three NAD(P)H+H^+^ (**Figure 2**). Violacein can be obtained from the fermentation of violacein-producing bacteria or by a Ruthenium-catalyzed chemical synthesis in organic solvents^40^. The pathway can be heterologously expressed in other bacteria^41^ and has been reconstituted *in vitro* from the five recombinantly produced enzymes^42,43^. The addition of catalase as a sixth enzyme improves reaction yield by supporting the intrinsic peroxidase activity of VioB^42^. More recently, the pathway became a model system for metabolic engineering in bacteria^39^, yeast^44,45^ or cell-free systems^46–48^.

**Figure 2.**
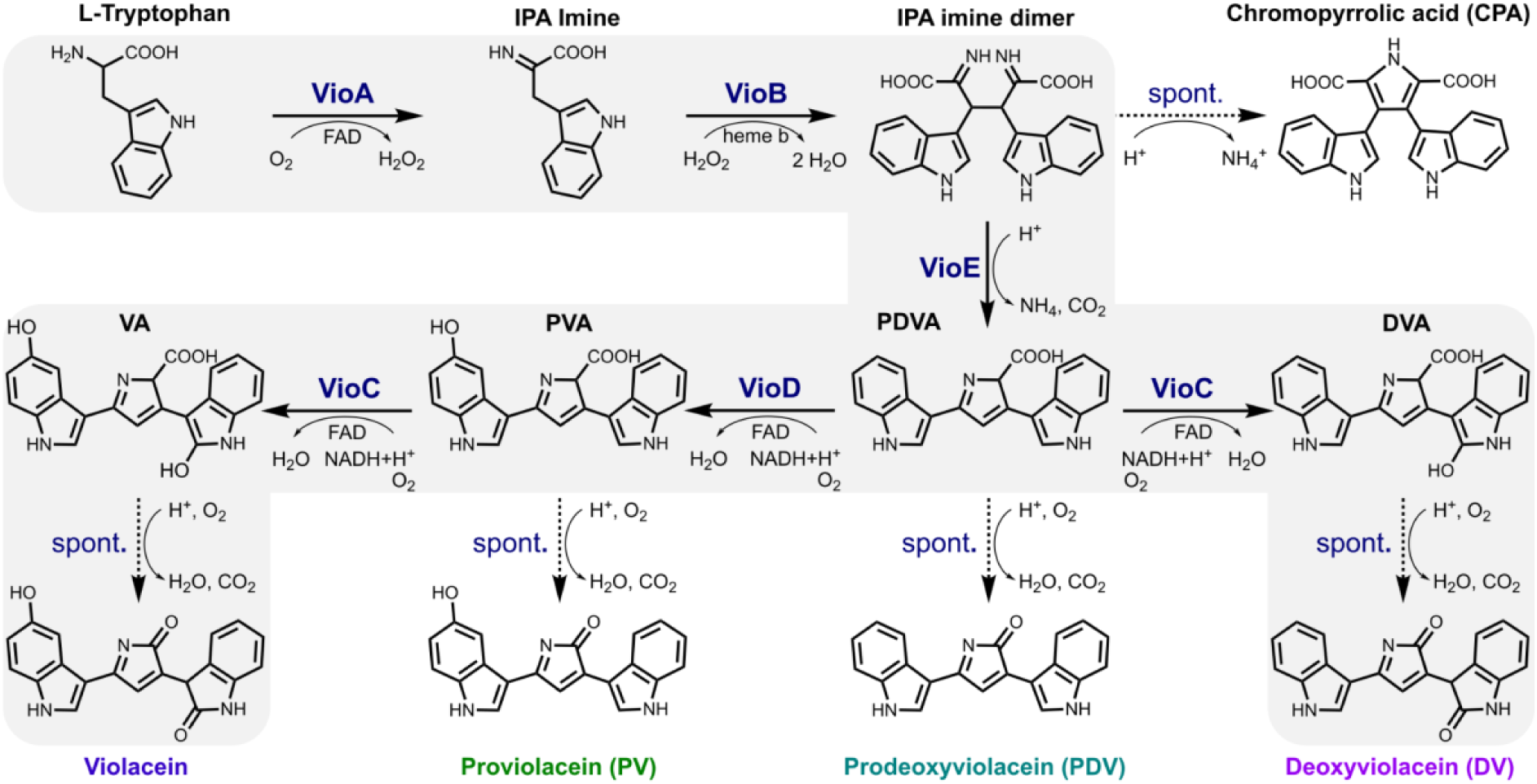
Violacein biosynthesis pathway. Violacein is formed from two molecules of L-tryptophan through five enzymatic steps catalyzed by VioA, B, E, D, and C, followed by a final non-enzymatic oxidative decarboxylation. Premature oxidation (or lack of an enzyme) leads to several pathway side products (CPA, PV, PDV). Deoxyviolacein is frequently observed as an alternative pathway product and assumed to have biological activities similar to violacein. Not shown: PV and PDV may (co-)react further to chromoviridans pigments^45^. DVA, deoxyviolaceinic acid; PDVA prodeoxyviolaceinic acid; PVA, proviolaceinic acid; VA, violaceinic acid.

We recombinantly produced all five enzymes VioA–E (**Figure 3A**) separately in *E. coli* and purified them using nickel affinity and size exclusion chromatography (**Table S1**). To ensure full cofactor loading, we supplemented flavin adenine dinucleotide (FAD) to the FAD-dependent proteins VioA, VioC and VioD during nickel affinity purification^42^. The heme-iron protein VioB could only be expressed in media supplemented with the heme precursors *δ*-aminolevulinic acid and ammonium iron sulfate^43^. Size, purity and oligomerization state of the five proteins were confirmed by SDS-PAGE (**Figure 3B**) and by analytical size exclusion chromatography coupled to multi-angle light scattering (SEC-MALS**, Figure S1**). As expected^49–51^, VioD was monomeric, whereas VioA and VioE appeared as homodimers. SEC-MALS and SEC coupled to small angle X-ray scattering (SEC-SAXS) showed both monomeric and dimeric VioB (**Figure S2B**), in line with a strong prediction for VioB dimers by AlphaFold^52^ (pTM score = 0.86, ipTM score = 0.89, Figure S2B). Initial preparations of VioC eluted as soluble high molecular weight assemblies sized between 0.4 to 1 MDa (**Figure S1C**). VioC expression at 16℃ and using purification buffers supplemented with 10% glycerol and 0.1% Tween20 resulted in mostly monomers (Figure S1). However, SEC-SAXS still identified higher molecular weight species (**Figure S2C**) and negative staining transmission electron microscopy (NS-TEM) revealed circular or semi-circular structures of about 16-17 nm diameter that would suggest a penta- or hexameric form (**Figure 3C**). AlphaFold predicts VioC homodimerization (pTM score=0.88, ipTM score = 0.86, **Figure S2D**) but not any higher-order multimers.

**Figure 3.**
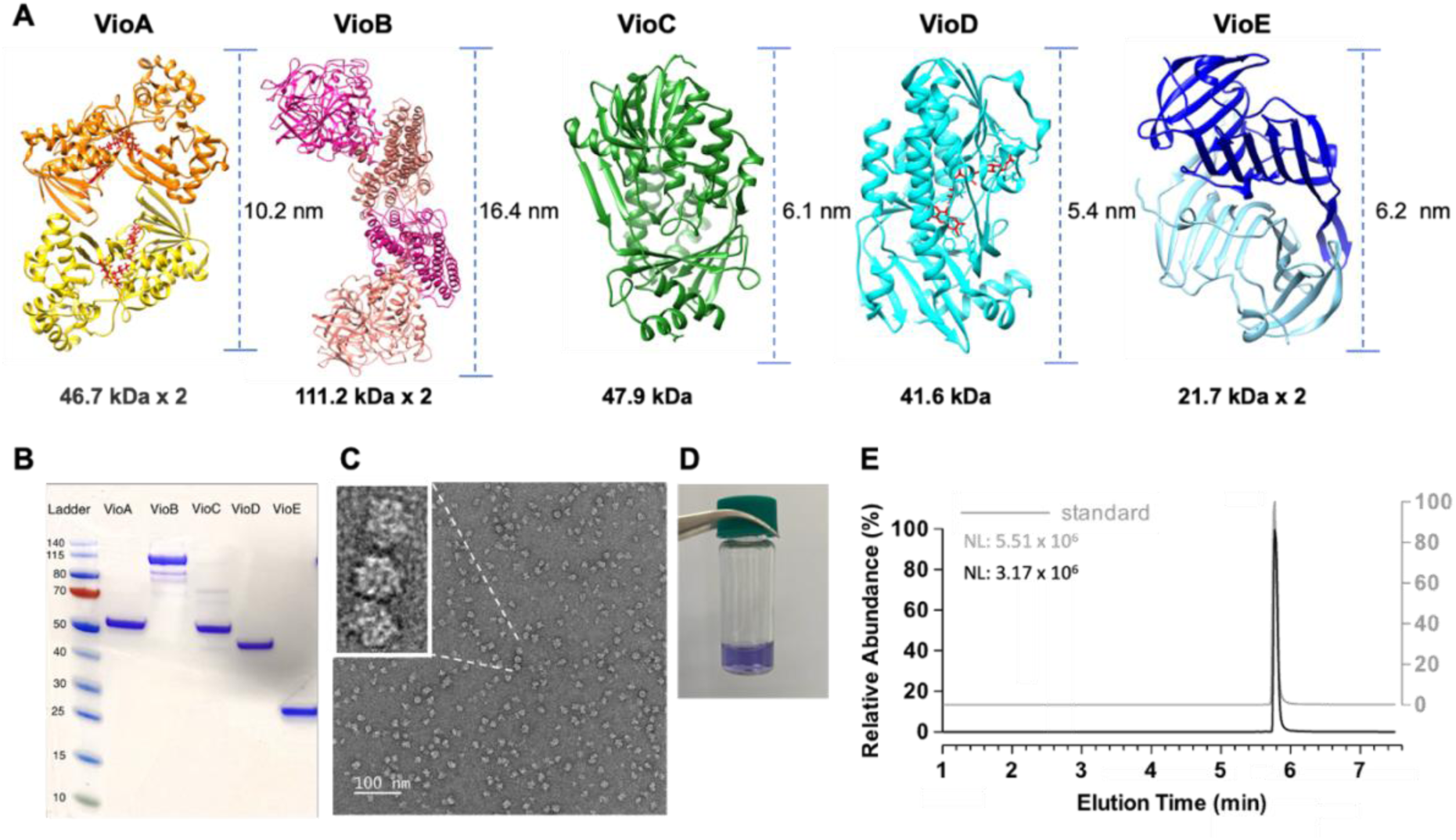
In vitro reconstruction of the violacein biosynthesis pathway. **A**, Three-dimensional structures of the natural violacein pathway proteins. VioA dimer (PDB: 5G3T), FAD (red); AlphaFold prediction of dimeric VioB (Q9S3V0); AlphaFold prediction of VioC (Q9S3U9); VioD (PDB: 3C4A), FAD (red); VioE dimer (PDB: 3BMZ). **B**, SDS-PAGE of the five purified enzymes. **C,** NS-TEM image of VioC indicating multimerization, magnification is 69k. **D**, Visual observation of violacein production in vitro. **E**, Violacein detection by UHPLC-MS/MS. The HPLC chromatogram was extracted using selected reaction monitoring mode in ESI (-) polarity.

We initiated violacein biosynthesis in solution by coincubating between 1 and 1.7 µM of each enzyme (VioA, B, C, D, and E, see methods for details) with cofactors (1 μM FAD, 5 mM NADPH) and 500 μM of the substrate L-tryptophan. We supplemented the reaction with catalase as a 6^th^ enzyme (50 U /200 µl reaction), to increase violacein yield^42^. The formation of the purple-colored violacein product was visible by eye (**Figure 3D**). Following methanol extraction, we quantified the product yield by ultra-high performance liquid chromatography tandem mass spectrometry (UHPLC–MS/MS). During the analysis, we filtered for product ions of violacein at 156 and 298 m/z in order to avoid confounding signals from pathway intermediates or side products (**Figure 3E**). Although there was variation between enzyme batches, we typically obtained product concentrations of approximately 20 to 45 µM, in line with or higher than those reported in the only previous publication of the fully reconstituted *in vitro* pathway reaction (25 µM)^43^. By comparison, higher yields (80 µM) were very recently reported for a cell-free pathway reconstitution with 10-fold higher substrate concentration and 24 h reaction times^48^.

### 2.2. Encapsulation and optimization of the pathway MIL system

ZIFs are currently the most widely used MOFs for protein encapsulation as they crystallize around proteins under conditions that are generally assumed non-denaturing^14,17^. We successfully encapsulated the VioA-E pathway enzymes into ZIF-8 (**Figure S3A-C**) but did not observe any pathway activity (**Figure S3D**). In fact, either of the two ZIF-8 components alone, Zn^2+^ or 2-methylimidazole, each abolished violacein production while the addition of pre-formed ZIF-8 crystals to free enzymes reduced product yield from 23 to 16 µM (**Figure S3E**). Individual single-enzyme encapsulation of only VioA, B, C, or D likewise blocked the pathway and only encapsulated VioE retained some activity (**Figure S3F**).

We therefore examined enzyme infiltration into MIL-101 (Fe)^35^ as an alternative. MIL-101 is characterized by a rigid architecture, high porosity, and excellent stability under moist conditions^35,53^. MIL-101 (Fe) is a good candidate for biological applications owing to its high biodegradability and low toxicity, including in mouse models^54^. However, MIL-101 features a very small pore size of only 3.5 nm which is why only small drug molecules could be encapsulated in this MOF^55,56^. We previously hierarchically engineered a MIL-101 structure with an expanded pore size ranging from 15 to 30 nm by controlled etching, and showed that this MIL-101 could be loaded with a model enzyme, catalase, that retained its catalytic activity^57^. For the present study we used this etched MIL-101 (Fe) (hereafter referred to as eMIL) to incorporate all components of the violacein biosynthesis pathway.

We synthesized MIL-101 (Fe) nanoparticles by the hydrothermal reaction of FeCl_3_x6H_2_O and terephthalic acid (H_2_BDC) in dimethylformamide (DMF) to obtain octahedral particles of 200 to 600 nm size. We then etched the MIL-101 nanoparticle by treatment with 100% acetic acid at 80℃ and varied the etching time to control the pore size^57^ (**Figure 4A**). Enzymes were infiltrated into eMIL particles by dropwise adding of a pre-mixed enzyme solution and stirring overnight at 4℃. Enzyme-loaded eMIL particles were collected by centrifugation and washed with buffer or water to remove any loosely absorbed protein (**Figure 4B**). We determined the efficiency of protein loading into the MOF indirectly from the protein concentration remaining in the supernatant and found that protein infiltration was most efficient (98-100%) with 90 min etched crystals (**Figure S4A, B)**.

**Figure 4.**
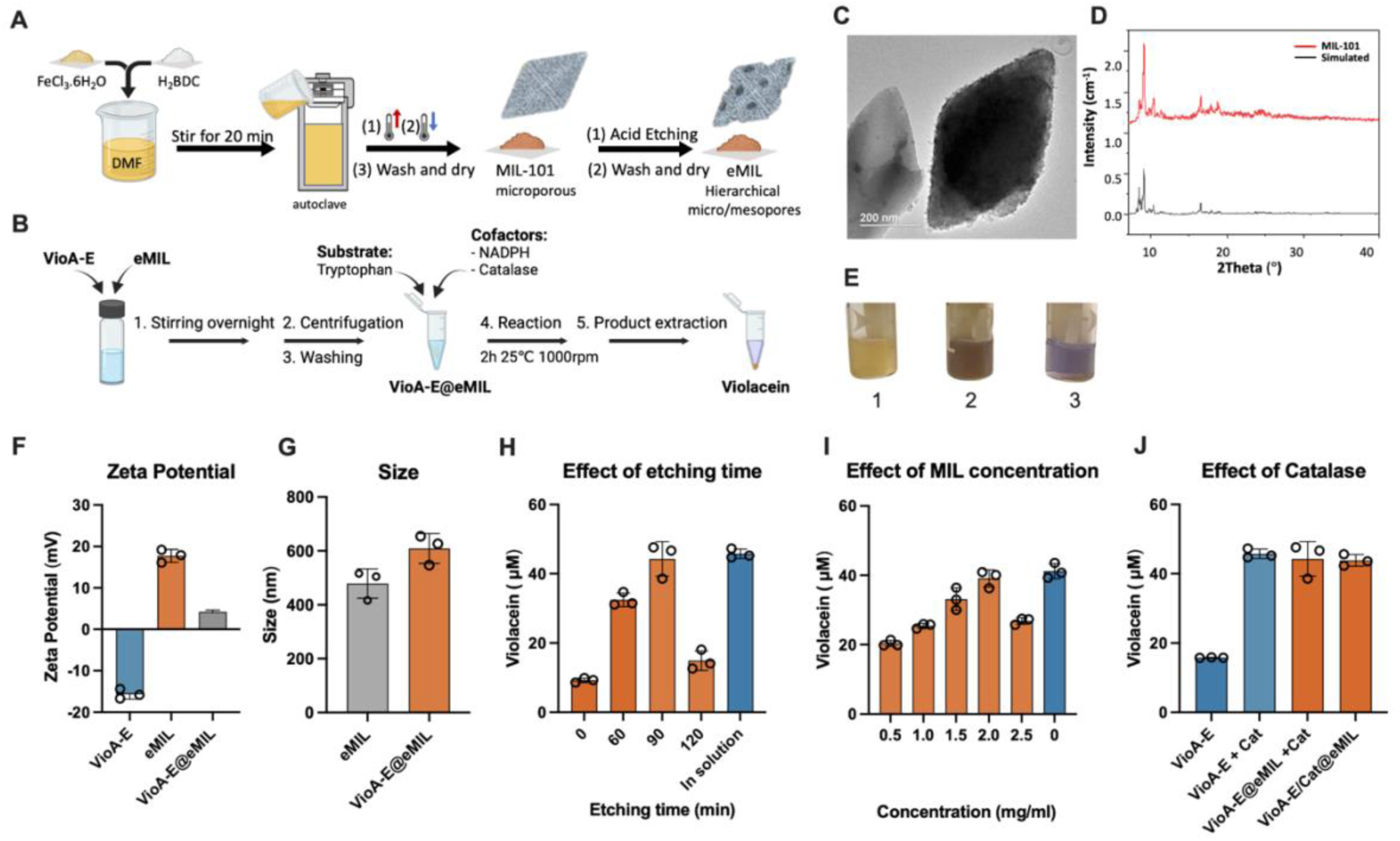
Variation and characterization of VioA-E@eMIL. **A**, Fabrication of MIL-101 followed by etching with acetic acid to form eMIL. **B**, Infiltration of proteins within eMIL. **C**, TEM images of eMIL particles, scale bar 200 nm (see also Figure S8). **D**, Powder XRD of eMIL. MIL-101-(90) indicates that the 90 min etching did not change the crystallinity of the eMIL. **E**, Visual evidence of violacein production: 1. eMIL alone, 2. VioA-E@eMIL after enzymatic reaction, 3. VioA-E reaction in solution. **F, G**, Zeta potential and size distribution of eMIL and VioA-E@eMIL. **H,** Comparison of violacein product yield for different etching times of MIL creating different pore sizes. **I,** Product yield for different eMIL / enzyme ratios. **J**, Effect of catalase on violacein production. Infiltration of five enzymes into eMIL with catalase supplied in the buffer (VioA-E@eMIL + Cat) or infiltration of all six enzymes into eMIL (VioA-E/Cat@eMIL) resulted in the same amount of violacein, on par with VioA-E and catalase acting in solution (VioA-E + Cat). **H, I, J,** Violacein was quantified by UHPLC-MS/MS. Error bars report the standard deviation of 3 replicates (data points shown).

Powder X-ray diffraction (XRD) confirmed that the crystallinity of the MOF did not change after 90 min etching (**Figure 4D**) and was also preserved after incubation in different buffers or cell culture medium (**Figure S5**). Etching also did not affect the decomposition temperature (400℃) in thermogravimetric analysis (TGA, **Figure S6**). Nitrogen adsorption / desorption at 77 K indicated that etching shifted the pore size distribution from 1.9 - 3 nm (MIL) to a broader range of peaks between 2 and more than 30 nm (eMIL) (**Figure S7**). The reduction in BET surface area^58^ from 700 to only 12 m^2^/g and the switch to a type II isotherm shape highlight a large-scale replacement of the original microporous structure with mesopores. Particles exhibited rough edges and tended to become more transparent in high-resolution transmission electron microscopy (TEM) images (**Figure 4C, Figure S8**). Etching thus created a wide range of pore sizes throughout the whole particle volume.

Infiltration of the pathway did not significantly alter TEM images (**Figure S8**) and did not change crystallinity in PXRD (**Figure S9**). The main decomposition temperature of BSA-infiltrated eMIL remained unchanged and the increase in residual mass independently confirmed the complete loading of BSA to 25% loading capacity (**Figure S6**). However, the zeta potential of pathway-loaded eMIL decreased after infiltration, suggesting that proteins were not only inside the crystal but that some protein molecules were protruding out of or adhering to the surface (**Figure 4F**). Dynamic light scattering showed a slight increase in mean particle size from between 400 and 520 nm for the empty eMIL to 550 to 650 nm after enzyme infiltration which may suggest pкуarticle swelling and/or the formation of a "protein corona" on the MIL surface (**Figure 4G**). However, confocal microscopy of eMIL clusters infiltrated with fluorescently labelled BSA revealed that proteins were homogenously distributed throughout the entire MOF volume without any accumulation at the surface (**Figure S10A**). We similarly examined the co-location of pathway enzymes by infiltrating VioA-E while labelling the largest enzyme VioB, the potentially oligomerizing VioC and the small VioE each with a different fluorescent dye. Despite the difference in size, confocal z-stacks again showed all three proteins co-distributed throughout particle volumes without noticeable surface accumulation although their relative intensity sometimes varied between particles (**Figure S10B**).

We therefore proceeded to incubate the pathway-loaded eMIL with the violacein reaction buffer containing substrate and co-factors. Violacein formation was readily observed with the MOF reaction emulsion turning dark brown (**Figure 4E**), however only after switching to monomeric VioC preparations (**Figure S11**). When separated by methanol extraction and centrifugation, the MOF itself had changed color from light yellow to brown while the supernatant had adopted the characteristic purple (Figure S4C). Violacein product yield, quantified as before by UHPLC-MS/MS, was indeed highest for the 90 min–etched particles (**Figure 4H)**. Notably, some pathway activity was still observed for unetched particles, which implies that lower quantities of all five proteins can also adhere to the unmodified MIL surface (Figure 4H).

We varied the concentration of eMIL in our infiltration reaction from 0.5 to 2.5 mg/ml, and found that violacein production was maximized at 2 mg/ml corresponding to a 5.5-fold excess of eMIL over protein (**Figure 4I, Figure S4D**). At this ratio, enzymes were again infiltrated to completion with loading efficiencies of 97% or higher. However, in terms of protein load these conditions represent less than half of the eMIL’s apparent maximal loading capacity of >50% protein : eMIL mass (Figure S4D, **S12**). The infiltration supernatant remained devoid of any protein in SDS-PAGE, even after concentrating it 5-fold (**Figure S13**), further confirming the high loading efficiency. For comparison, we also tested the infiltration of all enzymes and BSA individually (**Table S2**). Interestingly the protein mixture (96-98%) appeared more efficient in filling out the mesoporous material than most of the individual proteins (74-99%). Protein infiltration was also stable over time, as demonstrated by the only minor escape of fluorescently labelled BSA during repeated washes over a 24 h period (**Figure S14**). In summary, multiple lines of evidence demonstrated that all pathway participants were infiltrated into eMIL equally and to completion (Figure S4, S6, S12, S13) and distributed throughout the whole particle volume (Figure S10) without affecting eMIL stability (Figure S6, S8, S9).

Under the refined conditions, violacein production from the pathway-loaded eMIL (VioA-E@eMIL) was comparable to the 2 h end point yield obtained from the same amount of enzymes in solution (**Figure 4I**). Up to this point, we had supplied the supporting catalase enzyme as a "cofactor" with the reaction buffer (at 50 units per 200 µl reaction, corresponding to 0.2 µg). We tested whether catalase could be co-infiltrated as a sixth enzyme into the MOF. Indeed, MOF incorporation of bovine liver catalase (a tetramer of ∼250 kDa) together with the VioA–E proteins (VioA–E/Cat@eMIL) was successful and gave the same violacein yield as the reactions with catalase in solution (**Figure 4J)**. We concluded that the eMIL hosted all six enzymes in their active state.

### 2.3. Performance of the eMIL-packaged violacein pathway

Several studies have reported that tight MOF-encapsulation can stabilize certain proteins against heat denaturation^7^. In our case, the MOF-stabilizing effect was uncertain as the enzymes are lodged inside the large etched pores of eMIL and may to some extent also be associated with the outer eMIL surface. To test for thermostabilization, we subjected VioA–E/Cat in solution or @eMIL to a 10 min heat treatment before measuring reaction activity at room temperature. Activity of the MOF-associated pathway remained stable after treatment at 37℃ and 42℃ and still showed measurable activity after the 50℃ treatment (**Figure 5A**). Conversely, activity of the enzymes in solution decreased by 50% and 75% after treatment at 37℃ and 42℃, respectively, and no violacein was produced after treatment at 50℃. Thus, the protein infiltration into eMIL provides significant thermostabilization.

**Figure 5.**
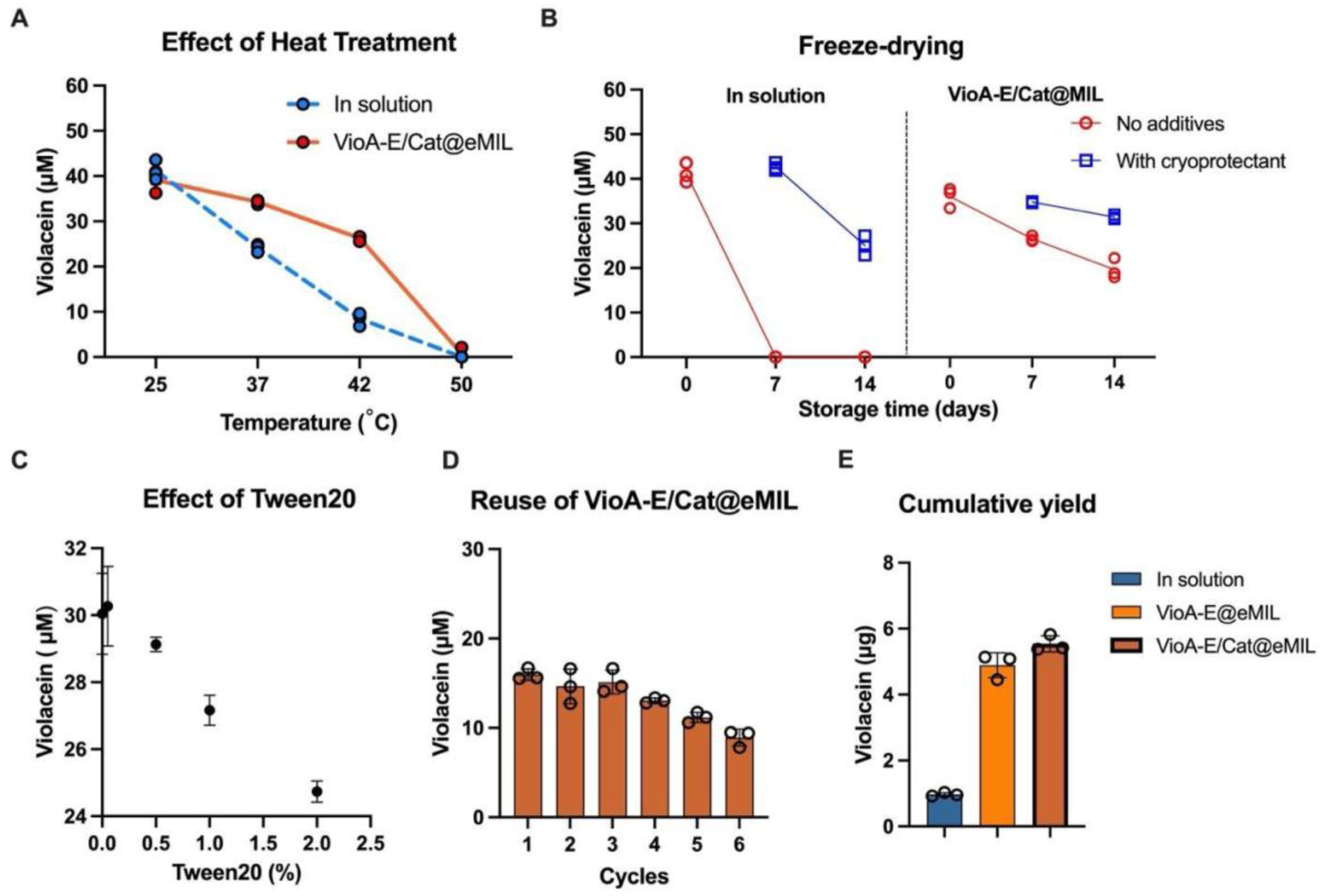
Performance of eMIL associated pathway. **A**, Violacein yield after a 10 min heat treatment of the pathway. **B**, Violacein production after lyophilization from solution (left) or from the enzyme-loaded eMIL (right), without (red) or with (blue) addition of cryoprotectant (5% mannitol, 5% trehalose), followed by storage and resolubilization. Day 0 corresponds to product yield prior to lyophilization. **C**, Tolerance of the Violacein pathway reaction against different concentrations of Tween20 (0.05%, 0.5%, 1%, 2%). **D,** Violacein pathway reuse over several cycles of biosynthesis. Violacein recovery from 100 µg of VioA-E/Cat@eMIL101 undergoing 6 cycles of 2h reactions. **E,** Cumulative violacein yield from all 6 cycles compared to the yield from a single reaction of the same amounts of enzymes and substrates in solution. All yields were quantified by UHPLC-MS/MS. Error bars in A-D report the standard deviation from three technical replicates (data points shown).

Protein transportation and handling is greatly facilitated by dry storage which, for individual proteins, is typically achieved through lyophilization. However, many proteins lose their activity after lyophilization, and cryo-protectants and conditions have to be optimized case by case^59^. MOFs have been reported to stabilize proteins under dry storage conditions^60^. We assessed the impact of lyophilization and subsequent -20℃ storage on VioA–E/Cat before or after MOF infiltration. Pathway lyophilization and freezing from solution without cryoprotectants completely abolished violacein production, whereas VioA–E/Cat@eMIL retained 50–75% of its initial activity after 7 and 14 days of storage (**Figure 5B)**. A common cryoprotectant mixture (5% trehalose, 5% mannitol) stabilized VioA–E/Cat when lyophilized from solution and further improved the protective effect of eMIL (Figure 5B). Therefore, eMIL, alone and in synergy with other compounds, acts as a cryoprotectant and facilitates the storage of the complete violacein pathway.

The perhaps most important advantage of enzyme immobilization or encapsulation is the straightforward separation of enzymes from small-molecule reaction products by filtration or centrifugation. This should enable the reuse of the biocatalysts in fresh reaction buffer, or their sustained activity in continuous flow reactors. We could indeed easily recover VioA–E/Cat@eMIL post-reaction by gentle centrifugation and washing. However, the violacein reaction product is hydrophobic and most of it remained associated with the VioA–E/Cat@eMIL. Methanol extraction^42,43^ denatured some or all proteins in solution but also within the eMIL, precluding their reuse. The nonionic surfactant Tween20 had recently been proposed as a green alternative to methanol for violacein extraction^61^. When screening reaction buffers with different Tween20 concentrations, we found that 0.05% Tween20 only marginally affected the violacein yield but allowed for the solubilization and extraction of most of the product under non-denaturing conditions (**Figure 5C**). We carried out successive rounds of violacein biosynthesis reactions in a buffer containing 0.05% Tween20, followed by centrifugation to recover the eMIL–protein complex, and addition of new buffer, substrate and cofactors for another reaction. Reaction yield in the supernatant decreased slowly over 6 cycles for both VioA-E@eMIL and VioA-E/Cat@eMIL but product formation remained high and the final cycle still reached more than 50% of the initial yield (**Figure 5D, Figure S15**). VioA–E@eMIL with and without catalase in the MOF produced similar total amounts of violacein, exceeding 5 µg over 6 cycles (**Figure 5E**). Overall, the 6-cycle violacein production was almost five times higher than the single cycle (∼1 µg) production achieved with the same batch and quantity of enzymes in solution (Figure 5E).

### 2.4. Reshaped pathway kinetics and reactant flows

We next assessed whether infiltration into eMIL altered the side products or kinetics of the pathway as compared to the enzymes in solution. We modified the UHPLC-MS/MS protocol to detect the four major potential pathway side products (CPA, DV, PDV, PV) and the central PDVA intermediate (see pathway diagram in Figure 2) by selective ion monitoring (SIM, **Table S3**) and validated the analytics of each of these compounds from reactions with incomplete pathways (**Figure S16**). However, as only a compound standard for violacein was available to us, the MS ion counts of the other compounds cannot be quantitatively compared to each other. All four side products and PDVA were detectable from a 2 h eMIL-infiltrated pathway reaction, whereas the solution reaction yielded markedly lower levels of DV and did not produce PDV (**Figure S17**).

In an effort to optimize our solution reaction conditions we first tested whether the five-fold increase in any one enzyme improved violacein yield (**Figure S18**). Balibar and Walsh had previously shown that VioB was rate limiting for the first three reactions (VioABE) but had not studied the full pathway ^42^. We observed that increases in VioB and, to a much lesser extent, in VioC or VioE, only enhanced the production of DV but not of violacein. Increasing VioA reduced violacein product, likely by diverting reaction flows towards a non-monitored side product ahead of the rate-limiting VioB. Increasing VioD eliminated DV but did not increase violacein production. We concluded that the violacein yield could not be improved by single enzyme increases.

To further probe the influence of relative enzyme concentration, we set up a new reaction containing 0.5 mg/ml total enzyme in equimolar ratio (1.7 µM each) with Tween20 included in both infiltration and reaction buffer to improve product solubilization and VioC function (see methods, Table S1). Rather than only analyzing the 2 h endpoint (**Figure 6A, B**), we additionally established pathway kinetics from reaction aliquots stopped at different time points (**Figure 6C, D**). Moreover, we performed experiments with five-fold diluted pathway concentrations for comparison (**Figure S19**). Violacein was again quantified by calibrated SRM (**Figure S20**) while all compounds were monitored by SIM.

**Figure 6.**
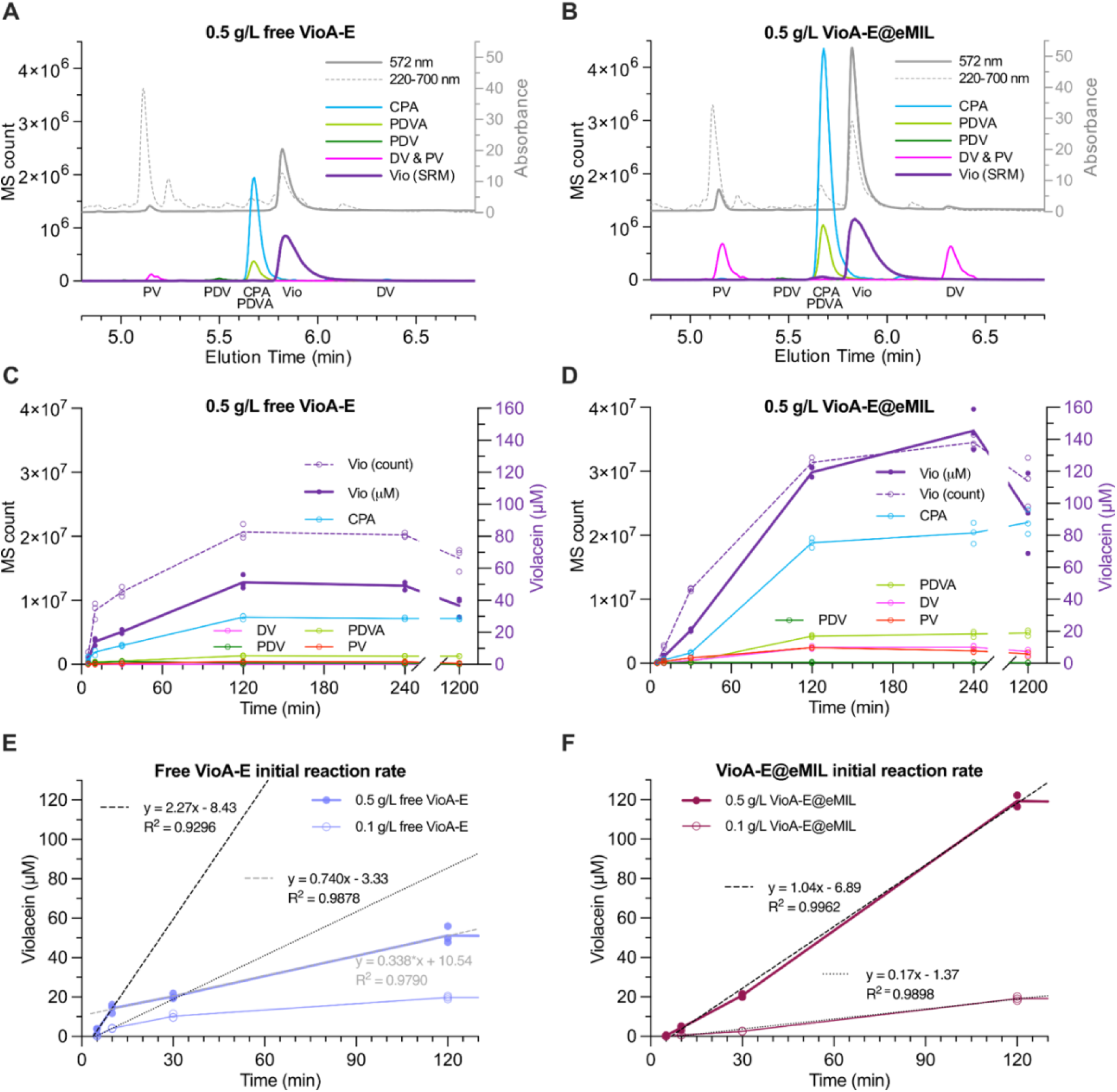
*In vitro* violacein pathway kinetics and side products. **A, B,** UHPLC-MS(/MS) elution profiles of VioA-E reactions at the 2 h time point for (**A**) free enzymes at 0.5 mg/ml total enzyme concentration (1.7 µM each) and (**B)** eMIL reaction with 0.5 mg/ml total enzyme infiltrated within 2 mg/ml eMIL. Reaction products were detected with SIM or SRM (Violacein only) and UV absorbance (gray, right axis). **C, D,** Time course of triplicated pathway reactions (**C**) of free enzymes and (**D)** of the equivalent eMIL reactions. Violacein concentration (right axis, purple line) was, as before, determined from SRM (MS/MS) calibrated against standard (Figure S20). The SIM count of violacein (left axis, purple dotted line) was recorded in parallel. Note that absolute CPA and PDVA signals benefit from the compounds’ negative charge. **E**, Violacein production from C and S19A replotted for the estimation of initial reaction rates of regular and diluted free enzymes reactions. Broken lines show linear regressions to only the 5 and 10 min triplicate points. Regression to a tentative secondary phase is shown in light gray. **F**, Violacein production from eMIL reactions replotted with linear regression to the 5 to 120 min time points. Reactions with five-fold diluted nanoreactors show the expected 5-fold reduction in initial reaction speed, in line with a prolonged steady state regime. See Figure S21 for the direct overlay of violacein production in the four conditions and S22 for a zoom in on lower abundance side products. The free pathway shows a higher initial rate of violacein production while the infiltrated pathway sustains the reaction for longer and to ∼threefold higher final yields. Abbreviations: chromopyrrolic acid (CPA), proviolacein (PV), prodeoxyviolacein (PDV), deoxyviolacein (DV), and violacein (Vio).

Enzymes in solution exhibited a rapid initial burst of violacein production, reaching 2.3 µM/min over the first 5 min (**Figure 6E**). In contrast (**Figure S21**), enzymes infiltrated in eMIL showed a prolonged lag phase (6.6 min *versus* 3.4 min for free enzymes) before maintaining a steady violacein production rate of 1.0 µM/min for at least 2 h (**Figure 6F**). The free reaction reached its peak of 51 ± 4.3 µM violacein after 2 h (Figure 6C), whereas violacein concentration from the VioA-E@eMIL nanoreactors kept increasing from 119 ± 4.0 µM at 2 h to a maximum of 145 ± 13 µM after 4 h (Figure 6D). Product precipitation impeded analysis of reaction yields after 4 h (Figure 6D). By calculating the ratio of initial reaction velocity to enzyme concentration, the apparent turnover number (k_cat_) for the full pathway was estimated to be ∼1.4 min^-1^ for enzymes in solution (diluted reaction ∼2.2 min^-1^) and ∼0.6 min^-1^ (diluted reaction ∼0.5 min^-1^) for eMIL-hosted enzymes (Figure 6E, F). Although violacein biosynthesis has not previously been studied at this level of detail, these rates fall within the range of other complex biosynthesis pathways for which *in vitro* turnover has been reported ^62–64^.

At and beyond the 2 h time point, the infiltrated pathway generated a larger amount of violacein than the free enzyme reaction but also increased concentrations of all detectable side products (Figure 6A, B**, Figure S22**). Notably, the pronounced accumulation of both DV and PV suggest limiting activity of VioD, which may lead to the saturation of VioC with the PDVA intermediate. Pathway output could therefore likely be further enhanced by a more complex rebalancing of enzyme concentrations. Importantly, the prolonged lag phase, reduced initial rate, and increased side product formation, each rule out substrate channeling^65,66^ as a mechanism behind the three-fold increase in final violacein yield. Overall, these results demonstrate that enzyme infiltration within eMIL fundamentally reshapes pathway kinetics with opportunities for optimization and mechanistic insight beyond conventional solution-based enzyme systems.

### 2.5. Intracellular pathway delivery

Violacein has been proposed as a potent anti-cancer agent. This activity has been ascribed to various mechanisms, including induction of cell death, inhibition of proliferation and migration, or hyperpolarization of the cell membrane cells^67–70^. In the present study, rather than delivering the violacein natural product, our goal was the intracellular delivery of the VioA-E plus catalase enzyme cascade for violacein synthesis *in situ*. Intracellular violacein production would rely on the availability of the tryptophan substrate and the NADPH cofactor within the cell. Tumor cells maintain higher levels of NADPH than non-tumor cells, because of their dependence on aerobic glycolysis (the Warburg effect)^71,72^. We hypothesized that the MOF-delivered violacein pathway may therefore also be more cytotoxic to cancer cells than to normal cells.

eMIL alone showed an acceptable cytotoxicity towards two cell lines with IC_50_ values of approximately 150 µg/ml (**Figure S23A**), in line with previously reported values^56^. As a proof-of-concept for the internalization of eMIL and delivery of cargo, we first infiltrated BSA labelled with either Alexa Fluor 647 or FITC into eMIL and incubated BSA@eMIL with HeLa cells for 16 h. The cells displayed overlapping bright red and green fluorescence in the cytoplasm (**Figure 7A)**, whereas cells did not take up free BSA (**Figure S24)**. We tested different incubation times by flow cytometry and found that, already after 4h, nearly all cells had taken up FITC-labelled BSA@eMIL (**Figure S25**). Intracellular BSA@eMIL manifested as black particles in the cytosol, outside of the nuclear envelope, in transmission electron microscopy (TEM) of fixed cell slices (**Figure 7B,C)**. White (empty) spots around some particles suggested the occasional splintering and displacement of the more rigid nanoparticles during the cross-sectional cutting with the microtome. Confocal microscopy, flow cytometry and TEM therefore collectively confirmed the intracellular localization and persistence of protein-loaded eMIL.

**Figure 7.**
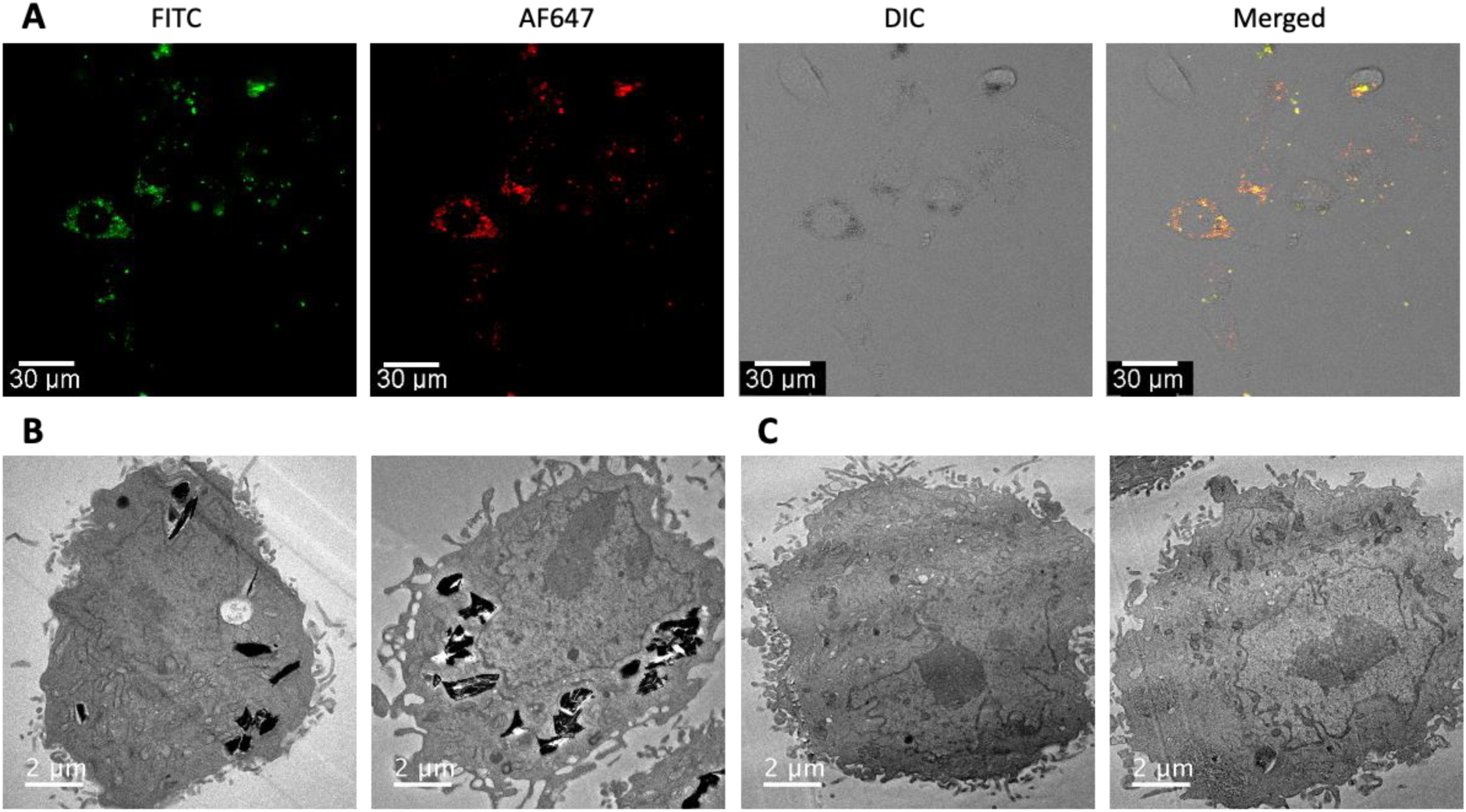
Internalization of BSA-infiltrated eMIL within HeLa cells. **A**, Confocal fluorescence microscopy of live cells after 12 h exposure to 50 µg/mL eMIL infiltrated with a mixture of FITC-labelled and Alexa fluor 647 (AF647)-labelled BSA, shown in green and red. DIC: differential interference contrast. **B**, Transmission electron microscopy (3.8k magnification) of cell slices fixed after 24 h exposure to 100 μg/mL eMIL-101 infiltrated with BSA. Nanoparticles remain largely intact and outside of the nucleus. **C**, Untreated control cells.

Although the violacein natural product has anti-cancer properties^68^, it is also toxic for normal cells^60^. We therefore next studied the toxicity of violacein itself, infiltrated in eMIL or given in solution, for a cancer-derived and a non-cancerous cell line (HeLa and 3T3, respectively). Violacein doses correlated with cell death, albeit HeLa cells tended to be more resistant to violacein than 3T3 cells (**Figure 8A**). In comparison with free violacein treatment, the packaging into eMIL (violacein@eMIL) increased toxicity in 3T3 cells but had less effect on HeLa cells (**Figure S23B, C**).

**Figure 8.**
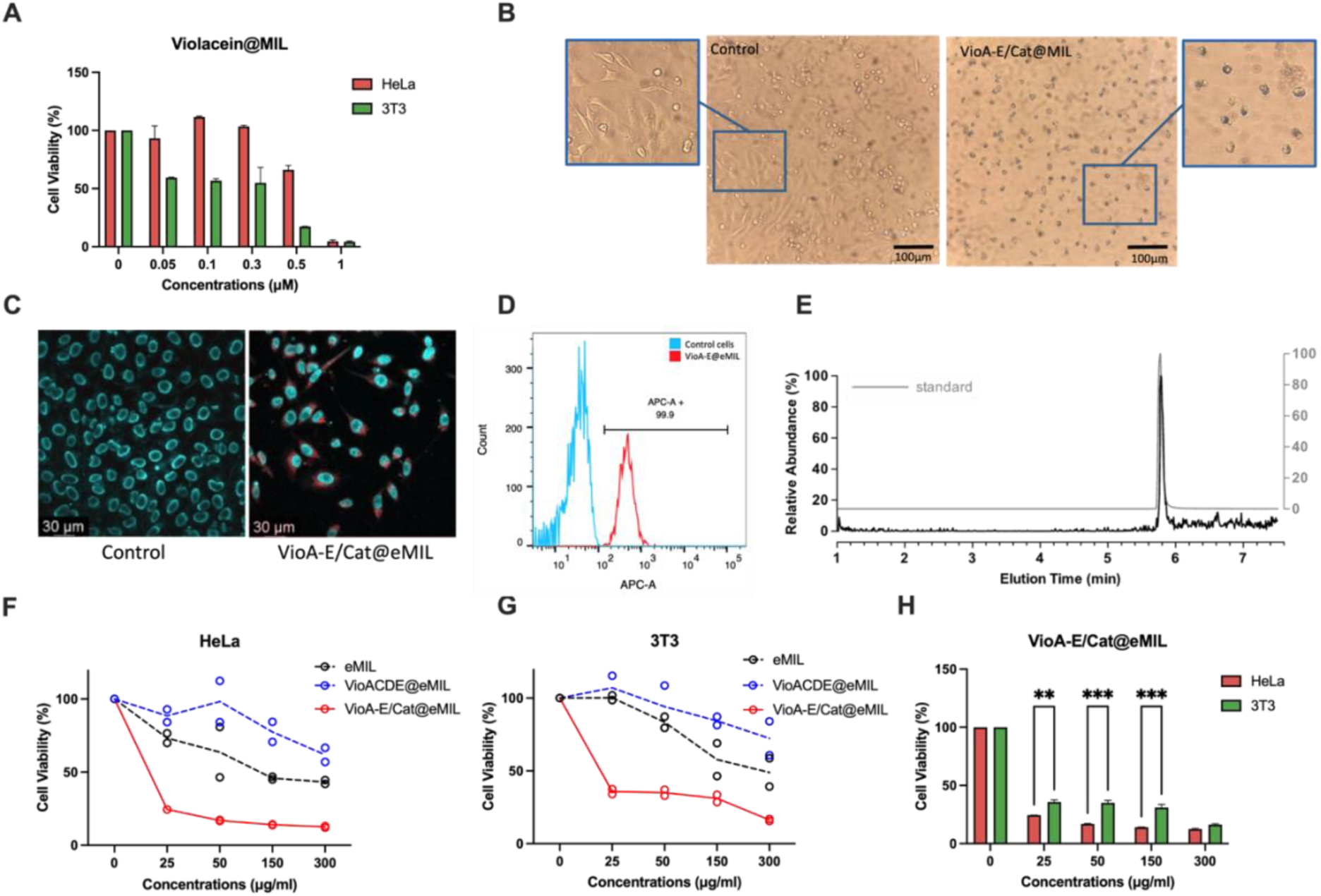
Intracellular pathway delivery. **A,** Cell viability of Hela and 3T3 cells at different concentrations of Violacein@MIL. **B,** Light microscopy of Hela cells treated with VioA-E/Cat@eMIL. Purple color indicates intracellular violacein production leading to apoptosis. Scale bar: 100µm. **C,** Confocal microscopy of HeLa cells incubated overnight with 50 µg/mL of eMIL (control) and VioA-E@eMIL. Nuclei were stained with Hoechst 33342 (cyan). Violacein produced intracellularly is visible in the Rhodamine channel (red). Scale bar: 30 µm **D**. Cells producing violacein were identified as APC-positive in flow cytometry analysis. **E**, UHPLC-MS/MS chromatogram detecting violacein extracted from HeLa cells after VioA-E/Cat@eMIL treatment. **F-G**, Cell viability of HeLa cells (**F**) and 3T3 (**G**) at different concentrations of VioA-E/Cat@eMIL, eMIL, and an incomplete-pathway control (VioACDE). **H**, Cell viability of HeLa and 3T3 cells at different concentrations of VioA-E@eMIL. Cell viability in A, F, G, H was assessed with the CCK assay and normalized to results from control cells without treatment. Error bars in A report the standard deviation from 3 technical replicates. The reported mass concentrations always refer to the pure eMIL not accounting for the added protein.

We then proceeded to deliver the full VioA–E/Cat@eMIL pathway into cells. Violacein production inside HeLa cells was evident from the purple color displayed after incubation with VioA–E/Cat@eMIL which we did not observe in cells incubated with unloaded eMIL or free enzymes (**Figure 8B**). Cell shapes pointed to wide-spread apoptosis occurring only in the pathway-treated cells (**Figure 8B**). The intrinsically fluorescent violacein stained the cytosol in confocal microscopy (**Figure 8C**) and turned nearly all (remaining) cells fluorescent in flow cytometry (**Figure 8D**). The natural product remained tightly associated with cell pellets when mixed with methanol or various detergents (**Figure S26**) but could be extracted with chloroform:methanol (2:1) and was identified by UHPLC-MS/MS (**Figure 8E**). Thus, eMIL infiltration delivered all enzymes in an active state into the cell, where they were functioning in concert as a multi-step natural product biosynthesis pathway.

Importantly, incubation of HeLa and 3T3 cells with VioA–E/Cat@eMIL for 24 h reduced cell viability substantially (**Figure 8F-G**), whereas cell viability was not affected by incubation with free enzymes (**Figure S23D**). To prove that toxicity was caused by intracellular violacein production, we excluded VioB from the eMIL-infiltrated enzymes resulting in a nanoreactor that cannot produce violacein (VioACDE/Cat@eMIL). Indeed, VioACDE/Cat@eMIL only showed moderate cytotoxicity at higher concentrations (**Figure 8F-G**), comparable to empty eMIL (compare also Figure S23A). VioA–E/Cat@eMIL reduced the viability more for HeLa cells than for the non-cancerous 3T3 cells, whereas violacein-loaded eMIL was more cytotoxic for 3T3 cells (**Figure 8H**). Enhanced cytotoxicity of the nanoreactor for cancer cells was further supported by the observation that ≥50 µg/ml VioA–E/Cat@eMIL also decreased the cell viability of the breast cancer cell line MCF-7 more than of HEK-293 (**Figure S23E**). The enhanced toxicity in cancer cells is in line with their higher content of the NADPH cofactor. However, metabolic reprogramming may also affect tryptophan availability^73^ and the transplanted pathway may behave differently in different types of cancer or healthy tissue. Collectively, these results demonstrated that eMIL infiltration enabled the intracellular delivery of a fully functional enzyme pathway, with its performance being sensitive to the cellular metabolism.

## 3. Conclusion

Herein, we demonstrated that a complex biosynthetic pathway of up to six enzymes can be reconstituted within hierarchically etched MIL-101 (eMIL). eMIL was able to host large quantities of enzymes with diverse sizes and charges, bypassing limitations of ZIF-8 encapsulation, which, by contrast, impaired enzyme activity. eMIL confinement altered pathway dynamics in unexpected ways: It increased lag phases and side products but, nevertheless, tripled overall yield. This suggests that the MOF microenvironment modulated enzyme behavior through complex inhibitory and protective effects, possibly involving pH shifts, substrate access, intermediate entrapment, or redox activity from Fe³⁺ centers^74^. These mechanisms warrant further investigation. Apart from outperforming free enzymes *in vitro,* nanoreactors stabilized the pathway system for storage and reusability. The eMIL’s high carrying capacity enabled simple, highly efficient and, based on current experiences, nearly indiscriminate protein loading. With “no protein left behind”, it was easy to quantify and change the nanoreactor protein composition by simply adjusting the enzyme mixture in the infiltration solution. Collectively, these properties identify eMIL nanoreactors as a robust platform for *in vitro* biosynthesis, in particular for smaller-scale applications.

Remarkably, eMIL enabled the intracellular delivery of the whole pathway into mammalian cells, where all enzymes behaved as a single functionally integrated protein system that produced violacein from endogenous substrates. The dependence on host cell metabolism likely contributed to the increased violacein production, and hence cytotoxicity, in cancer cell lines compared to normal cells. We cannot exclude that some intracellularly produced violacein may diffuse out and affect neighboring cells, although we have no evidence for it. If such effects were significant, they could be advantageous in therapeutic contexts. eMIL particles are unlikely to release their infiltrated proteins and appear stable in the cytosol, thus creating a protein-protective but highly substrate and product diffusible synthetic reaction compartment inside the cell.

To our knowledge, violacein-producing nanoreactors represent the first example of protein-based “pathway transplantation” and the most complex multi-protein system delivered into living cells to date. Given the biocompatibility of the original MIL-101 in mice^54^, eMIL-based multi-enzyme delivery seems well positioned for transient metabolic engineering aimed at therapeutic applications. The spikes and throughs of traditional systemic drug administration often lead to risks of toxicity as well as subtherapeutic exposure, while exposing the entire organism to potential side effects. In contrast, pathway-infiltrated eMILs introduce the concept of protein *system* therapies, where therapeutic molecules may be synthesized locally and maintained at more stable concentrations only in the tissues where they are needed. Metabolic rewiring through eMIL nanoreactors also presents a compelling alternative to gene therapy, avoiding complex and risky *in vivo* genetic manipulation. Importantly, our work exemplifies that delivering active pathways—rather than final products—enables environment-responsive, “smart” therapies. Additionally, practical advantages such as enhanced stability and dry storage address some of the issues that complicate access to conventional biologics. We conclude that pathway–MOF nanoreactors create new opportunities for the engineering and deployment of biocatalytic systems, with broad implications for both technical applications and next-generation precision medicine.

## 4. Experimental Methods

### Materials

Flavin adenine dinucleotide disodium salt hydrate, β-nicotinamide adenine dinucleotide 2′-phosphate reduced tetrasodium salt hydrate, bovine liver catalase (catalogue C100), δ-aminolevulinic acid, ammonium iron-(II)-sulfate and violacein standard were purchased from Sigma. HeLa (CCL-2) and MCF7 (HTB-22) cell lines were obtained from ATCC. HEK-293 was obtained from Merck (85120602). The cell cytotoxicity assay kit (ab112118) was purchased from Abcam. Lab-Tek II 4 Chambers cover glass used for cellular uptake imaging experiments was purchased from Nunc.

### Protein expression and purification

Amino acid sequences for *Chromobacterium Violaceum* ATCC 12472 VioA to E genes were obtained from Uniprot: VioA (Q9S3V1), VioB (Q9S3V0), VioC (Q9S3U9), VioD (Q9S3U8), VioE (A0A202BAH3) and reverse-translated and codon-optimized for the expression in *E. coli* using an in-house Python script based on DNAChisel^76^. A SpyTag^77^, 3C protease cleavage site and 8xHis tag were added C-terminally. Annotated sequences are provided in supplemental materials. DNA constructs were gene-synthesized by Twist Bioscience (US) in our pJEx411c expression vector with Kanamycin resistance, lacI, T7P/lacO, and a RBS insulator cassette (BCD2) that improves translation initiation^78^. Plasmids were transformed into chemically competent *E. coli* BL21(DE3) cells (Invitrogen) according to the manufacturer’s protocol. Starter cultures were inoculated from single colonies and grown shaking at 37°C in Terrific Broth (TB) media (IBI Scientific) overnight. 1L TB production cultures supplemented with 50 mg/L kanamycin were inoculated 1:100 (v:v) and incubated at 37°C and 225 rpm shaking. Protein expression was induced with 0.1 mM isopropyl-β-D-thio-galactoside (IPTG, final concentration) around an optical density (OD) of 0.6-0.8 and left shaking overnight at 25°C. For the expression of heme-containing VioB, 1 mM δ-aminolevulinic acid and 40 μM ammonium iron-(II)-sulfate were added (final concentrations). VioC was grown at 37°C in TB, induced with 0.1 mM IPTG, and incubated overnight at 16°C. Following a small-scale expression screen, VioD yield and purity was in later batches improved by induction with 1 mM IPTG and expression over night at 20 °C. Cells were harvested by centrifugation at 6000 g for 10 min and resuspended in binding buffer (25mM Tris pH 7.4, 500mM NaCl, 10 mM Imidazole, 10% glycerol and 1mM DTT) supplemented with SigmaFast protease inhibitor cocktail, EDTA-free, 1 tablet/100ml (Sigma Aldrich) and Benzonase (10 U/ml), and lysed in a Cell Disruptor (Constant Systems, UK) at 15 kPsi. The lysate was clarified by centrifugation at 50,000 g for 50 min at 4°C and filtered over Miracloth (Merck). Proteins were then captured on Ni-NTA resin in gravity columns for 1h at 4°C and washed five times with the binding buffer. FAD-dependent enzymes, VioA, VioD, VioE were further incubated with 4 mM FAD for 1h at 4°C and washed five times. Proteins were eluted with increasing concentrations of 50, 250 and 500 mM imidazole. After concentration in spin-concentrators (Millipore), all proteins were subsequently subjected to size-exclusion chromatography in 20 mM Tris-HCl pH 7.5, 500 mM NaCl, 10% glycerol, 1mM DTT on a Superdex200 16/600 column (Cytivia) using an Åkta FPLC (GE Healthcare). All buffers for VioC purifications were supplemented with 0.1% Tween20 which removed higher order oligomerization observed with the original purification. Protein purity was analyzed by SDS-PAGE (12% Bis-Tris, MES buffer, Invitrogen) and Fairbanks staining^79^. Selected SEC peaks were concentrated using Amicon Ultra 15 mL centrifugal filters (30 kDa MWCO), aliquoted, flash-frozen in liquid Nitrogen and stored at -80°C.

### SEC-MALS and SEC-SAXS

Size exclusion chromatography combined with multiangle light scattering was performed in 20 mM HEPES, pH 7.5, 200 mM NaCl, 1 mM TCEP, 0.01% (w/v) NaN_3_ on a Superdex200 10/300 column (Cytiva) at a flow rate of 0.75 ml/min using an Agilent HPLC setup coupled to a Dawn Heleos II and OptiLab T-rEx (Wyatt, U.S.A.) instrument. Protein was injected at a concentration of 2.5 mg/ml. Analysis was performed with the Astra 7.7 software (Wyatt Technology). Protein UV (280 nm) extinction coefficients were later calculated in Astra 8.4 from experiments on an updated SEC-MALS setup with Agilent 1260 Infinity II G7165A MWD UV detector, Dawn Neon MALS, and OptiLab differential refractive index (dRI) detector (Wyatt / Waters, U.S.A). The dRI signal provides an accurate sequence-independent protein (mass) concentration which allows for the determination of a (mass-concentration) UV extinction coefficient within the protein’s SEC peak. Multiplication with the (sequence-defined monomer) molar mass yielded an experimentally defined molar extinction coefficient. Owing to FAD or heme prosthetic groups in all but the VioE enzyme, these deviated from the sequence-derived extinction coefficients we had originally used for protein concentration measurements^80^. All protein concentrations reported in this study are based on the experimental extinction coefficients (Table **S1**).

SEC-SAXS experiments were performed at the SWING beamline (SOLEIL, Saint-Aubin, France), using the Biology Laboratory 1 (proposal number 20220373) as reported previously^81^. However, the buffer used was 20 mM Tris, 300 mM NaCl. Data were processed and analysed using SWING’s on-site FOXTROT software and PRIMUS of the ATSAS software package.

### In vitro enzymatic reaction and quantification

Violacein pathway reactions were carried out in 200 µl violacein buffer (20 mM Tris-HCl pH 8, 100 mM KCl) containing 0.36 mg/ml total protein (1.2 μM VioA, 1.4 µM VioB, 1 µM VioC, µM VioD, 1.7 µM VioE), 500 μM L-tryptophan, 1 μM FAD, 5 mM NADPH and 50 units of catalase (0.1 µg). The reaction mixture was incubated at room temperature for 2 h shaking in a thermomixer at 1000 rpm. 400 µl methanol were then added to each sample, vortexed and centrifuged at 13,000 rpm for 10 min to precipitate proteins and, where applicable, MOF. Violacein was quantified on a Thermo Vanguish Flex UHPLC system coupled to a Thermo TSQ Altis Plus Triple Quadrupole Mass Spectrometer after separation on a Zorbax Eclipse Plus C18 column (2.1 x 150 mm, 3.5 μm particle size, Agilent) with water and acetonitrile supplemented with 0.1% formic acid as the mobile phases. 5 μL of the methanol/water supernatant were injected and subjected to a gradient of 5 - 98% acetonitrile over 9 min run time at 0.5 ml/min flow rate. The MS was operated in negative electrospray ionization mode using the following parameters: capillary voltage=3500 V, sheath gas=35 Arb, Aux gas=14 Arb, ion transfer tube temperature=325°C, vaporizer temperature=350°C. Selection reaction monitoring (SRM) with two transitions of violacein (156.9 and 298.1 m/z) were used to specifically detect violacein. Quantification was based on the area of two transitions (TIC) on HPLC, then area counts were transformed into concentration using a calibration curve prepared from commercially available standard (Sigma-Aldrich; V9389-1MG).

### Encapsulation of enzymes within ZIF-8

VioA-E@ZIF-8 was synthesized by mixing 5μM of each violacein enzyme with 2-methylimidazole (final concentration 2.25 M). The zinc nitrate solution (final concentration 0.5 M) was slowly added while stirring for 20 min. The resulting product was collected by centrifugation and washed three times with a buffer (20mM Tris pH 8, 100 mM KCl) to remove any residues. The protein concentration remaining in the supernatant was measured by Bradford assay calibrated against BSA and used for calculating loading capacity and encapsulation efficiency.

### Synthesis and characterization of etched MIL-101

MIL-101 (Fe) was synthesized according to previous literature^57^. Briefly, 2.45 mmol Fe_2_Cl_3_ x 6H_2_O and 1.24 mmol H_2_BDC were mixed in 15 mL DMF and stirred for 5 min. Next, the mixture was heated in an oven at 110°C for 20 h without stirring. The mixture was centrifuged and washed with DMF and dried at 110°C. Etching of MIL-101 was performed by stirring for 20 minutes with glacial acetic acid at 80°C for between 90 and 120 min. Finally, products were centrifuged and washed several times with DMF and then distilled water.

The zeta potential of eMIL was measured on a Malvern Zetasizer Nano ZS at 25°C at pH 7.3 in aqueous solutions. PXRD patterns were recorded using Cu-K*α* radiation (1.54056 Å) on a Bruker D2 Phaser. TEM images were obtained using FEI Tecnai 12 microscope operating at 120 kV. For visualization by TEM, samples were prepared by dropping the solution on a copper grid 300 mesh (Electron Microscopy Sciences, LC 300-Cu).

The thermal stability of MIL-101, eMIL and eMIL loaded with BSA was evaluated by thermogravimetric analysis (TGA) using a TGA 5500 (TA Instruments). All samples were heated from 25 to 1000°C at the rate of 10 °C/min, using N_2_ as the protective gas. N_2_ gas adsorption measurements were performed using a Micromeritics ASAP 2020 instrument after sample degassing at 150°C for 12 h. The sorption measurement was maintained at 77 K under liquid nitrogen. The surface area was measured by the Brunauer-Emmett-Teller (BET) method.

### Infiltration of enzymes within eMIL

Infiltrations were performed in 20 mM Tris-HCl pH 8, 100 mM KCl with 0.36 mg/ml total enzyme and 2 mg/ml eMIL, optionally supplemented with 25 U/100 µl of catalase. In detail, a three-fold concentrated enzyme mix was slowly added while stirring to a 3 mg/ml eMIL suspension. The solution was stirred overnight at 4°C in a glass vial. eMIL was collected by centrifugation and washed three times with water. All supernatants were collected and subjected to Bradford assay to calculate the loading capacity and infiltration efficiency.

### Protein release from eMIL

For the release profile of protein infiltrated in eMIL, BSA conjugated with fluorescein isothiocyanate (BSA-FITC) was used as a model protein. BSA-FITC was infiltrated into eMIL as before (0.5 mg/ml BSA-FITC + 2 mg/ml eMIL) and washed and diluted into PBS. Triplicates of 2 ml release reactions containing 1 mg/ml BSA-FITC@eMIL (eMIL-only concentration) in PBS, pH 7.0, were incubated at 37 °C under gentle shaking. At predetermined time intervals, samples were centrifuged 5 min at 10,000×g, the supernatant was collected and fluorescence intensity was measured (excitation/emission: 495/525 nm) to determine the concentration of released BSA-FITC. Free BSA-FITC (not encapsulated) of the same final concentration (0.25 mg/ml) was used as a reference corresponding to 100% release. Cumulative release was calculated as a percentage of the total fluorescence signal from the free BSA-FITC control.

### Pathway reusability and stability

Reusability of VioA-E@eMIL was evaluated by measuring Violacein yield after repeated usage of the same eMIL pellet. Tween20 was added to the reaction at a final concentration of 0.05%. Pathway-loaded eMIL was recovered by centrifugation and then suspended in a fresh reaction solution (including substrate and cofactors). The reaction supernatants from every cycle were collected and separately quantified by UHPLC-MS/MS.

A Labconco instrument was used for lyophilization of VioA-E@eMIL or enzymes in solution according to the manufacturer’s instructions. Samples were flash frozen in liquid nitrogen and placed overnight in the lyophilizer until dry and then kept at -20°C. Where indicated, VioA-E@eMIL and free enzyme solutions were lyophilized in a cryoprotectant buffer (20mM Tris pH 8, 100 mM KCl, 5% trehalose, 5% mannitol). 2 h enzymatic reactions were performed after a 10 min heat treatment at 25, 37 or 50°C. Violacein was extracted with two volumes methanol and centrifugation at 13,000 rpm for 10 min. As before, 5 μL of the methanol extract was injected for UHPLC-MS/MS analysis.

### Microscopy

For the imaging of particle uptake into cells, HeLa cells were seeded at 20,000 cells/well in 4-chamber slides and maintained overnight in complete DMEM media at 5% CO_2_ and 37°C. 10 µg/ml of BSA-AF647@eMIL and FITC-BSA@eMIL or 2.5 µg/ml of free BSA-AF647 and FITC-BSA were incubated with the cells for 4 h at 37°C. Hoechst dye was added to the cells to stain the nucleus and incubated in the dark for 5 min. Cells were imaged using the same microscope (Leica SP8) using a 40x objective lens. following a standard protocol. The uptake of the free and eMIL-loaded BSA was detected using two laser channels (Hoechst @405/460, AF647 @650/668). Cells with no treatment were used as a control to set the intensity and gain. For the visualization of intracellular violacein production, HeLa cells were seeded into 96 well plates to 8,000 cells/well overnight at 37°C and 5% CO_2_. Cells were then treated with 300 µg/ml eMIL and enzymes@eMIL and incubated for 24 h. Cells were imaged using an inverted laboratory microscope (Leica DMIL LED) through a Leica DFC3000 G objective at 20x magnification.

For the fluorescence detection of intracellularly produced violacein, HeLa cells were seeded at 20,000 cells/well in 4-chamber slides and maintained overnight in complete DMEM media at 5% CO_2_ and 37°C. 50 µg/ml of VioA-E/Cat@eMIL and eMIL alone were incubated with the cells overnight and stained with Hoechst dye as explained earlier. The cells were imaged using the same microscope (Leica SP8) using a 40x objective lens. The uptake of the free and VioA-E/Cat@eMIL was detected using two laser channels (Hoechst @405/460, Violacin@Rhb(546/590).

### Particle microscopy

0.5 mg/ml FITC-BSA (Sigma Aldrich, A9771) or BSA-AF647 (ThermoFisher, A34785) in violacein buffer (20 mM Tris-HCl pH 8, 100 mM KCl) was left stirring with 2 mg/ml eMIL overnight, collected by centrifugation, washed three times and supernatants were subjected to Bradford assays for the determination of infiltration efficiency as before. For direct particle imaging, FITC-BSA@eMIL was diluted to 0.125 mg/ml (note that concentrations always refer to eMIL alone even for protein-loaded particles), sonicated and 20 µl applied to glass slides with coverslips and imaged on a Leica SP8 confocal microscope with a 100x/1.40 oil immersion objective, excitation at 491 nm, emission at 500-550 nm, 1 Au pinhole, at 400 Hz bidirectional scanning and 100 step Z-stack with 0.17 µm step size.

For triple labeled VioA-E@eMIL imaging, enzyme aliquots were concentrated into labelling buffer (20 mM HEPES pH 7.0, 150 mM NaCl, 0.02% Tween20, 0.02% sodium azide, 0.04 mM TCEP) with Amicon Ultra 0.5 ml centrifugal filters and then labelled as follows: VioB with Alexa Fluor 488 C5-maelimide (A10254), VioC with Alexa Fluor 594 C5-maleimide (A10256), VioE with Alexa Fluor 647 C2-maleimide (A20347). 50 or 100 µl reactions with with 50 µM (VioB) or 100 µM (VioC, VioE) enzyme and two-fold molar excess of dye were incubated at room temperature for 2h in light-protected microcentrifuge tubes. Excess dye was removed using Zeba 0.5 ml spin desalting columns (7 kD MWCO) and protein and dye concentrations were measured on a nanodrop instrument. The resulting labelling efficiencies were between 0.5 (VioC, VioE) and 0.8 (VioB) per protein. Labelled (VioBCE) and unlabelled (VioAD) enzymes were combined into a 100 µl infiltration reaction in violacein buffer with 2 mg/ml eMIL and 0.5 mg/ml enzyme mix (equimolar 1.7µM each) and incubated in a foil-wrapped 4 ml glass vial and left stirring (200 r.p.m.) overnight at 4°C. The infiltrated eMIL was then collected and washed twice with violacein buffer and resuspended in MilliQ water at mg/ml of eMIL (not considering protein load). A 10 µl drop of the resuspended eMIL was added to a glass slide and covered with a glass coverslip. Imaging was performed on the same setup as described above but with separate excitations at 488 (laser strength 3.8), 561 (7.2) or 633 nm (6.8) and emissions at 500–550 nm, 570–620 nm, or 650–700 nm in line-sequential mode and a 50 step Z-stack was collected at 0.14 µm step size.

### Transmission electron microscopy (TEM)

HeLa cells were seeded in 6-well-plates (100,000 cells/well) and cultured overnight at 37°C in 5% CO_2_. The following day, the cells were treated with 100 μg/mL of BSA@eMIL for 24 h, then washed three times with PBS and fixed overnight a in solution of 2.5% glutaraldehyde, and 2% paraformaldehyde in 0.1 M sodium cacodylate buffer. Subsequently, the cells were washed three times with sodium cacodylate buffer. Post-fixation was performed using 1% osmium tetroxide and 1.5% potassium ferrocyanide for 1 h in the dark. Following post-fixation, the cells were washed three times with water and stained with 2% osmium tetroxide in distilled H_2_O for 30 minutes in the dark. After three additional washes with water, the cells were stained with 1% uranyl acetate overnight at 4°C in the dark. Cell samples were dehydrated using increasing concentrations of absolute ethanol and subsequently embedded in epoxy resin. Subsequently, 70‒100 nm sections were cut with a diamond knife using a microtome and placed on TEM grids. The samples were post-stained with lead citrate for 2 minutes, washed with distilled H_2_O, and visualized using a Titan ST instrument (FEI company) and analyzed using Digital Micrograph software.

For pure particle images, approximately 1 mg of MIL-101, eMIL or protein-loaded eMIL was dispersed in 1 ml of ethanol with the help of sonication. The suspension was drop-cast onto a 300-mesh copper grid and allowed to dry under ambient conditions. High-resolution transmission electron microscopy (HR-TEM) was performed using the Titan ST instrument equipped with OneView camera, operated at an accelerating voltage of 300 kV.

### Cytotoxicity / viability assay

Cell viability was quantified with the Cells Cytotoxicity Kit (ab112118, Abcam) following the manufacturer’s protocol. Briefly, cells were seeded at 8,000 cells/well into a 96-well plate and incubated overnight at 37°C and 5% CO_2_. Cells were treated with different concentrations of eMIL, VioA-E@eMIL, free violacein, and violacein@eMIL in 200 µl total volume. Cells without treatment were used as a control, and all treatments were performed in triplicates. After 24 h, medium was exchanged for 100 μl of fresh DMEM with 10 μl of CCK-8 and incubated for 3-4 h. Absorbance at 570 nm and 605 nm was then measured on a microplate spectrophotometer (Thermo Scientific™ Varioskan™ LUX multimode microplate reader). The 570/605 absorbance ratio was calculated after subtracting the average absorbance value of wells without cells. Wells with cells but without any treatment were used as a control and their mean value was considered as 100 % viable. The viability of treated cells was calculated as (Corrected Mean Absorbance of treated cells)/(Corrected Mean Absorbance of control) ×100%.

### Flow cytometry analysis

HeLa cells were seeded into 6-well-plates (50,000 cells/well). After reaching confluency, cells were incubated with 50 µg/mL VioA-E@eMIL for 24 h, washed with phosphate-buffered saline (PBS) and collected. Violacein was detected using the APC channel (at 633/670 nm) or the PE-Texas Red A channel (at 561/585nm) on a BD LSRFortessa flow cytometer equipped with BD FACSDiva (BD Biosciences) software.

To study the cellular uptake kinetics of nanoparticles, HeLa cells were incubated with with 50 µg/mL eMIL infiltrated with FITC-labelled BSA (FITC-BSA@eMIL) or 12.5 µg/mL free FITC-labelled BSA for durations of 4, 12, or 24 h. Cells were washed to eliminate any non-internalized MILs, and analysed on the same BD LSRFortessa flow cytometer.

### Pathway kinetics

VioA-E infiltrations were performed as before but with an equimolar enzyme mixture (1.7 µM each VioA-E) giving 0.5 mg/ml total protein mixed with 2 mg/ml eMIL in 20 mM Tris-HCl pH 8, 100 mM KCl and supplemented with 0.05% Tween20. Kinetics experiments were performed in the same buffer (including Tween20) with 0.1 or 0.5 mg/ml total enzyme (equimolar 0.34 or 1.7 µM each VioA to VioE). The equivalent VioA-E@eMIL reactions were prepared by resuspending the infiltrated eMIL to either 2 mg/ml or 0.4 mg/ml final concentration (as before, eMIL concentrations do not consider added protein content). As infiltration efficiencies were consistently near 100%, we did not correct for any protein loss. 200 µl reactions were performed in triplicates in 96-well plates and were started by combining enzyme (/eMIL) solutions with a premixed stock of substrate and cofactors. At each reaction time point (5’, 10’, 30’, 2h, 4h, 20h), 30 µL were withdrawn, mixed with 60 µl methanol, vortexed and then kept on ice for later processing. Samples were centrifuged, transferred into HPLC vials with glass insets and subjected to the UHPLC separation as described above. The SRM violacein detection was multiplexed with SIM filters for all other compounds as well as violacein (Table S3). Each SIM target was monitored within a +/- 0.5 Da mass window and peak areas were integrated within a predefined retention time window (typically ±0.1 min). Proviolacein and deoxyviolacein were captured by the same SIM filter but could be distinguished by retention time (Figure S12 D, E). For VioA-E@eMIL, enzyme-limited steady state conditions were indicated by the ∼5-fold decrease in rate (0.17 µM/min) for the 5-fold diluted pathway (Figure 6F) albeit the PDVA intermediate only stabilized after 2 h. For the free reactions, linearity could not be assured at our low time resolution and the five-fold diluted pathway showed a three-fold lower initial rate (0.74 µM/min) (Figure 6E).

### Quantification and statistical analysis

All statistical analysis and visualisations were conducted with the Prism v.9.0 software (GraphPad). Data are reported as means ± standard deviation. Statistical significance was calculated by two-way ANOVA and Tukey’s multiple comparisons test: \**P* < 0.05, \*\**P* < 0.01, \*\*\**P* < 0.001, and \*\*\*\**P* < 0.0001. Chemical structures were produced with Chemdraw.

## Acknowledgements

We acknowledge SOLEIL for provision of synchrotron radiation facilities and would like to thank J. Perez and A. Thureau for assistance in using the beamline SWING. Experimental research was supported by the Bioscience Core Lab, the Imaging and Characterization Core Lab as well as the Analytical Core Lab at KAUST. Figure 1 was created by Heno Hwang, scientific illustrator at KAUST.

