## Supplemental Information for "Hierarchically engineered multi-enzyme nanoreactors for *in vitro* drug biosynthesis and pathway transplantation into cells"

**Table S1**. Protein characteristics

| ID[1] | Full enzyme name (cofactor) | E.C.[2] | $M_r$[3] kDa | pI | $\epsilon_{280}$[4] $(M\ cm)^{-1}$ | Oligo state | PDB | Uniprot | References[5] for oligo state |
|---|---|---|---|---|---|---|---|---|---|
| VioA (as1001) | L-tryptophan oxidase (FAD) | 1.4.3.23 | 46.7 | 7.2 | 102500 | dimer | 5G3T 5ZBD 6ESD | Q9S3V1 | Füller 2016 Yamaguchi 2018 Lai 2021, p. 201 this study (Fig. S1) |
| VioB (as1023) | 2-imino-3-(indol-3-yl) propanoate dimerase (heme b) | 1.21.98.- 1.11.1.6 | 111.2 | 6.0 | 183900 | dimer / monomer | - | Q9S3V0 | this study (Fig. S1) |
| VioC (as1003) | Violacein synthase (FAD) | 1.14.13.224 | 47.9 | 9.2 | (67300) | monomer / variable assemblies | - | Q9S3U9 | this study (Fig. S1) |
| VioD (as1004) | Protodeoxyviolaceinate monooxygenase (FAD) | 1.14.13.217 | 41.6 | 6.5 | 74300 | monomer | 3C4A | Q9S3U8 | Ran 2015 this study (Fig. S1) |
| VioE (as1005) | Protodeoxyviolaceinate synthase | - | 21.7 | 7.2 | 53800 | dimer | 3BMZ 2ZF3 | - | Ryan 2008 Hirano 2008 Asamizu 2007 this study (Fig. S1) |
| Cat | Catalase | 1.11.1.6 | 61.3 | 5.4 | | tetramer | 6PM7 | P00432 | Herskovits 1969 |

**Notes:** [1] Lab-internal unique construct IDs in brackets. [2] Enzyme classification number; [3] Monomer molecular weight; pI is the theoretical value (sequence-based) except for the well-documented experimental value for Catalase. Experimental values may differ substantially from sequence-based isoelectric point estimates; [4] Molar extinction coefficients at 280 nm per protein monomer (including spyTag and His tag) determined from differential refractive index (dRI) and UV absorbance of the main peak in SEC-MALS experiments. The VioC value could not be measured owing to Tween20 interference and was calculated from the sum of sequence-based and FAD extinction coefficients. [5]

**Table S2**. Infiltration of individual proteins into MIL-101 (constant at 2 mg/ml).

| **Protein** | **Infiltration mg/ml** | **Concentration in supernatant, mg/ml** | **Loading efficiency, %** | **Loading capacity, %** |
|---|---|---|---|---|
| VioA | 0.36 | 0.003 ± 0.001 | 99 ± 0.3 | 18 |
| VioB | 0.41 | 0.025 ± 0.001 | 94 ± 0.2 | 19 |
| VioC | 0.28 | 0.073 ± 0.015 | 74 ± 5.4 | 10 |
| VioD | 0.31 | 0.031 ± 0.005 | 90 ± 1.6 | 14 |
| VioE | 0.50 | 0.016 ± 0.001 | 97 ± 0.2 | 24 |
| Cat | 0.50 | 0.051 ± 0.005 | 90 ± 1.0 | 22 |
| BSA* | 0.50 | 0.044 ± 0.003 | 91 ± 0.6 | 23 |
| VioA-E* | 0.36 | 0.007 ± 0.002 | 98 ± 0.6 | 18 |
| VioA-E/Cat* | 0.36 | 0.016 ± 0.011 | 96 ± 3.1 | 17 |

Concentrations determined by Bradford assays calibrated against BSA (for individual enzymes) or against the actual enzyme mixture (VioA-E[/Cat]); Loading capacity = 100% massProtein/masseMIL; *Measured in a separate experiment

**Table S3**. UHPLC-MS/MS Detection parameters

| Name | Detection mode | [M-1] Da | Formula | Retention time, min | Validation Pathway |
|---|---|---|---|---|---|
| Chromopyrrolic acid | SIM | 384.4 | $C_{22}H_{15}N_3O_4$ | 5.67 | VioAB |
| Protodeoxyviolaceinic acid | SIM | 340.3 | $C_{21}H_{15}N_3O_2$ | 5.70 | VioABE |
| Prodeoxyviolacein | SIM | 310.3 | $C_{20}H_{13}N_3O$ | 5.38 | VioABE |
| Proviolacein | SIM | 326.3 | $C_{20}H_{13}N_3O_2$ | 5.08 | VioABED |
| Deoxyviolacein | SIM | 326.3 | $C_{20}H_{13}N_3O_2$ | 6.30 | VioABEC |
| Violacein | SRM | 342.3, [156.9, 298.1] | $C_{20}H_{13}N_3O_3$ | 5.80 | VioABEDC |

See methods for details. In brief, compounds were separated on a C18 column with a water : acetonitrile gradient from 5 - 95% (each with 0.1% formic acid) over a 10 min total run time at 0.5 mL/min. Intermediates and side products were detected by selected ion monitoring (SIM) mode at the given m/z (negative ion) while violacein was detected by selective reaction monitoring (SRM).

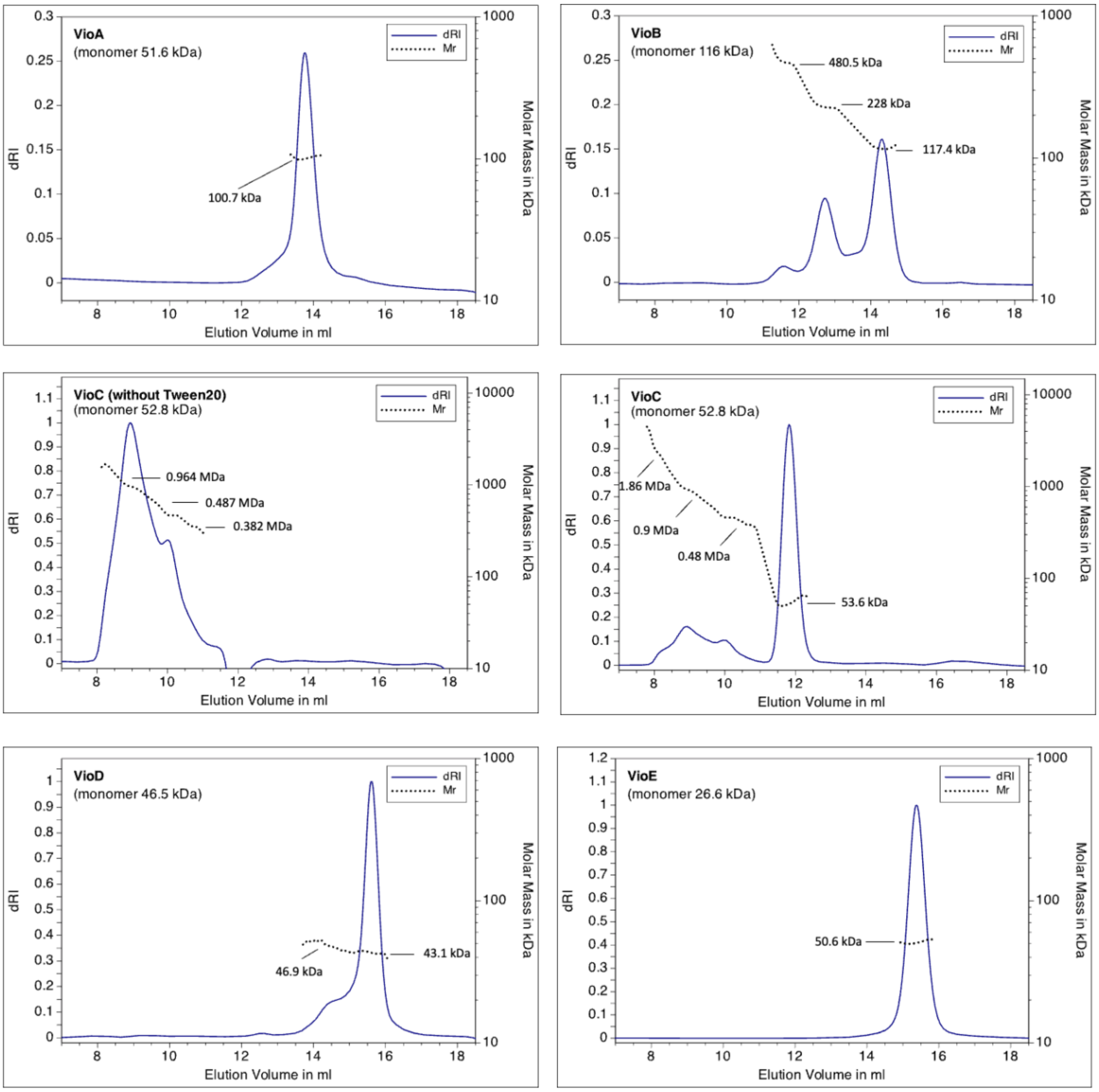


**Figure S1**. Oligomerization state as characterized by Size Exclusion Chromatography coupled to Multiangle Light Scattering (SEC-MALS). The expected molar mass for the monomer state given below each protein name corresponds to the full-length construct which always included SpyTag, 3C protease cleavage site and a C-terminal 8xHis tag. The molar mass estimated per elution point is superimposed onto the differential refractive index trace. $M_r$ value annotations were calculated from the wider peak area. Two independent SEC-MALS results are provided for VioC: initial preparations gave higher order oligomers of 0.4 – 1 MDa. Later purifications with Tween20 added during purification (but not in the SEC-MALS running buffer) yielded mostly monomeric enzyme. See text for further details.

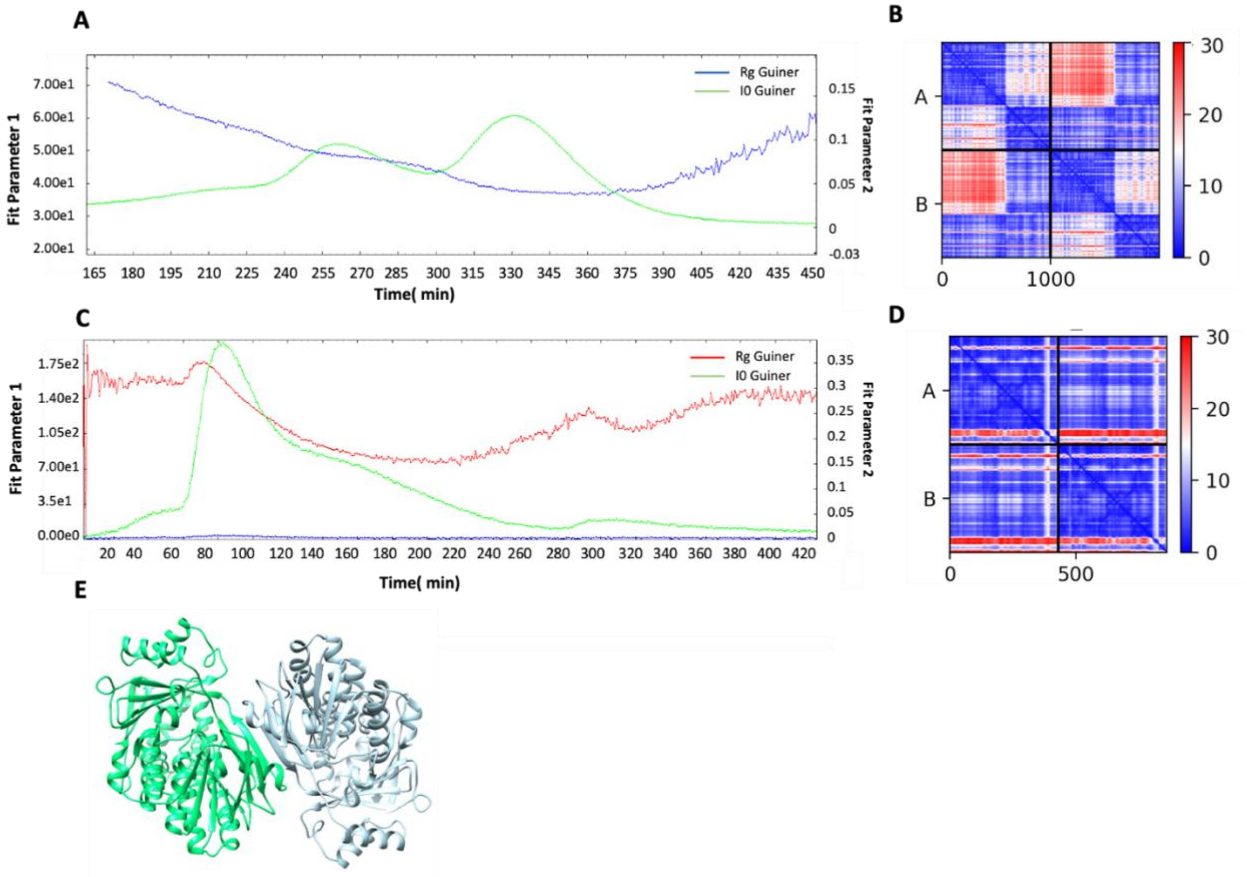


**Figure S2**. Additional analysis of oligomerization state of VioB and VioC proteins. **A**, Radius of gyration plot for VioB from SEC-SAXS. **B**, Predicted alignment error (PAE) graph of AlphaFold 2 structure of VioB dimer. pLDDT=92.18, pTM=0.86. **C**, Radius of gyration plot for VioC from SEC-SAXS. **D**, Predicted alignment error (PAE) graph of AlphaFold structure of VioC dimer. pLDDT=86.60, pTM=0.88. **E**, Corresponding AlphaFold prediction of dimeric VioC (Q9S3U9).

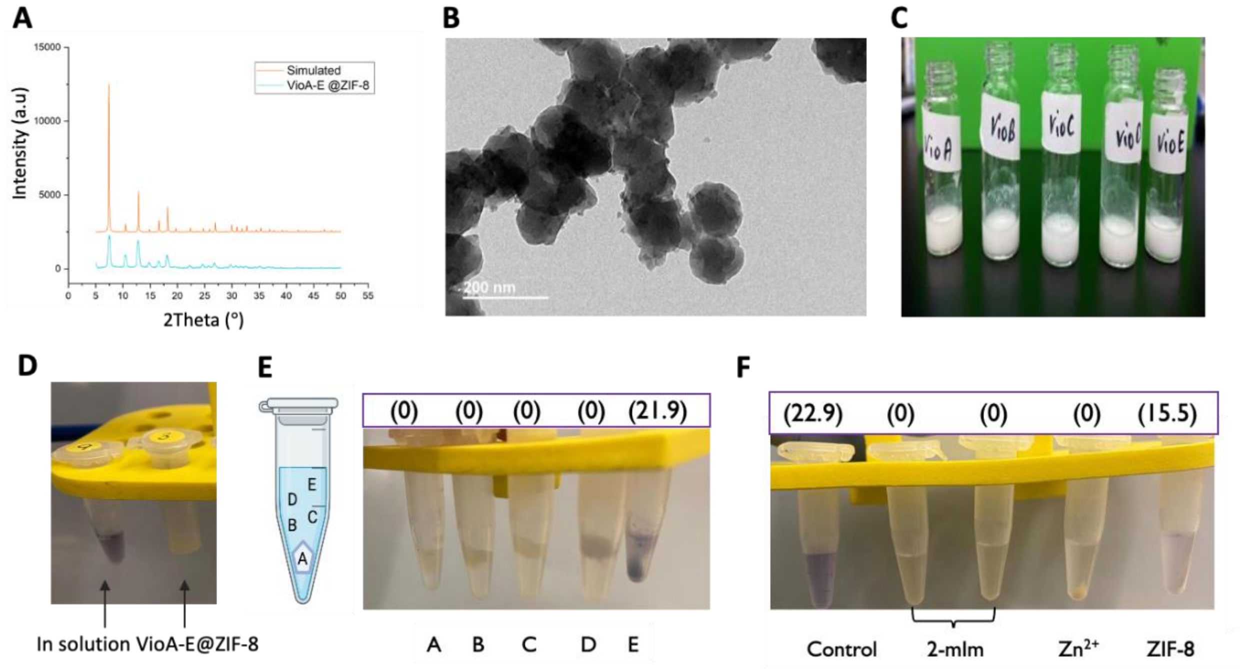


**Figure S3.** Encapsulation of the violacein pathway within ZIF-8. **A**. Powder X-ray diffraction (XRD) patterns of VioA-E enzymes encapsulated within ZIF-8. **B**. TEM image of ZIF-8 encapsulated VioA-E (VioA-E@ZIF-8). **C.** Formation of opaque solution confirmed successful encapsulation of VioA-E enzymes within ZIF-8. **D**. No violacein formation was observed with VioA-E@ZIF-8 compared to in solution enzymes. **E**. Visual observation of violacein production when only one enzyme was encapsulated within ZIF-8, while all others were added in solution. The Violacein concentration in µM as determined by HPCL-MS/MS is given above each reaction. **F**. Violacein yield after treatment of VioA-E enzymes with ZIF-8 components: Control, Enzymes in solution; 2-mlm, 2-methylimidazole (2.5M) for 1 min (left) and for 30 min (right); $Zn^{2+}$, Zinc nitrate (0.05M) for 30 min; ZIF-8, pre-formed ZIF-8 crystals; Violacein yields in µM as determined by HPLC-MS/MS are given in brackets above tubes.

**A**

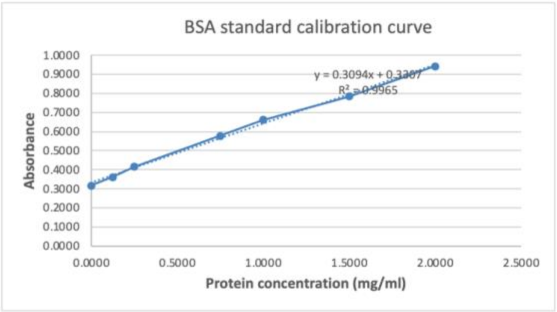


**B**

| Etching time (min) | Loading effiency (%) |
|---|---|
| 0 | 85 |
| 60 | 87 |
| 90 | 97 |
| 120 | 84 |

**C**

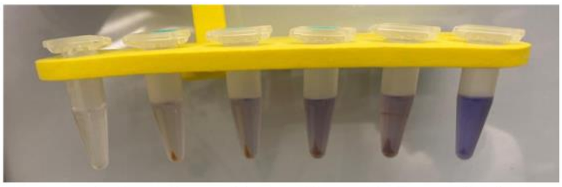


**D**

| [eMIL] mg/ml | Loading effiency (%) | Loading capacity (%) |
|---|---|---|
| 0.5 | 77 | 55 |
| 1 | 98 | 35 |
| 1.5 | 99 | 24 |
| 2 | 100 | 18 |
| 2.5 | 95 | 14 |

**Figure S4**. Optimization of VioA-E@eMIL infiltration. **A**, Bradford calibration curve using BSA. **B**, Etching time optimization: MIL etched for 0, 60, 90 and 120 min was loaded by mixing 2 mg/ml eMIL and 0.36 mg/ml enzyme mix and loading efficiency determined by Bradford assays in the supernatant. **C**, loading ratio optimization: visual observation of violacein production with different concentrations of eMIL (0.5 - 2.5 mg/ml) infiltrated with the same 0.36 mg/ml enzyme mix. **D**, Protein loading efficiency and loading capacity (protein : eMIL mass) for reactions in C as determined by Bradford assays in the supernatant.

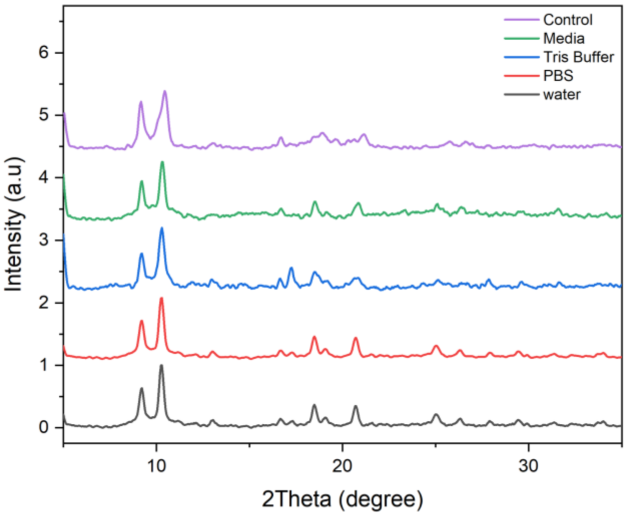


**Figure S5**. Preservation of eMIL-101 crystallinity in different solvents. eMIL were dissolved in water, Tris buffer, PBS or cell media for 24 h, then dried for analysis. The PXRD pattern of treated eMIL remains unchanged from that of the untreated control.

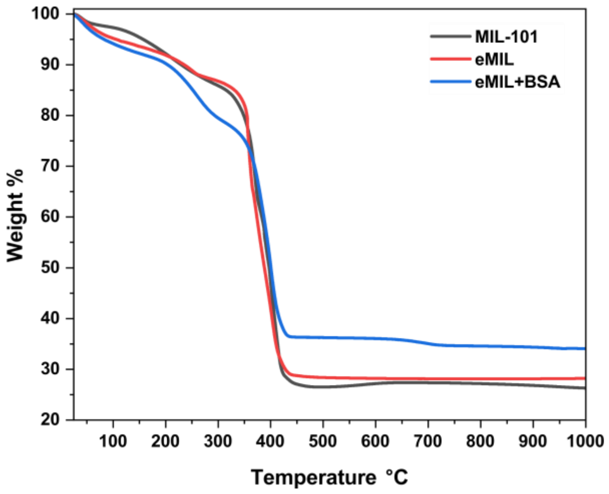


**Figure S6**. Thermogravimetric analysis. TGA was performed on unetched MIL-101, etched MIL-101 (eMIL) and eMIL infiltrated with BSA (0.5 mg/ml protein infiltrated into 2 mg/ml eMIL). MIL decomposition around 400 °C remains unaffected by etching and protein loading. The 25% (massProtein/masseMIL) protein load is reflected by a 25% increase in residual mass.

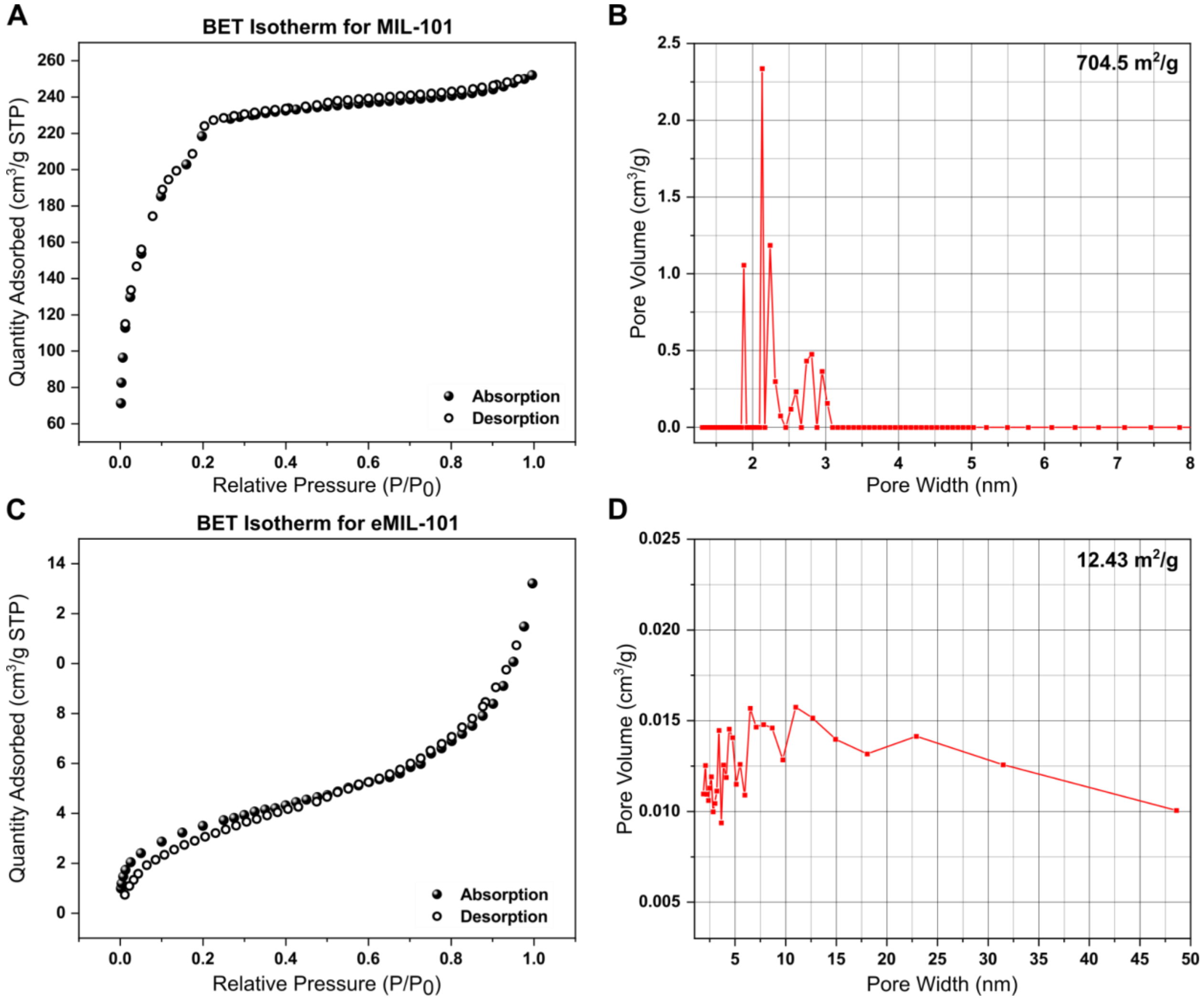


**Figure S7**. Assessment of surface area and porosity by N2 adsorption/desorption at 77 K. **A**, BET isotherms of unetched MIL indicative of type 1 with microporous attributes. **B**, Pore size distribution calculated using the DFT model of the BET isotherm in A. **C**, BET isotherms of (etched) eMIL-101. **D**, Pore size distribution for eMIL-101 calculated from B. All materials were degassed at 423 K (150 °C).

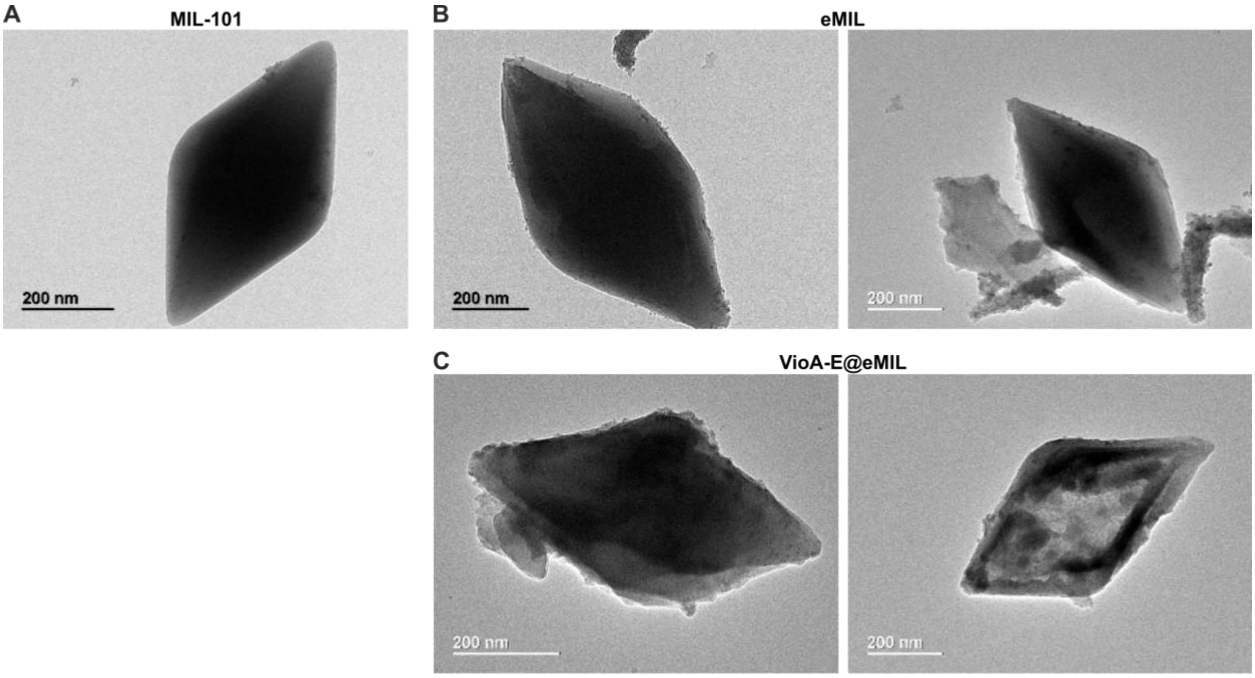


**Figure S8**. Representative TEM images. **A**, MIL-101 (before etching). **B**, etched eMIL. **C**, eMIL after infiltration with VioA-E pathway proteins.

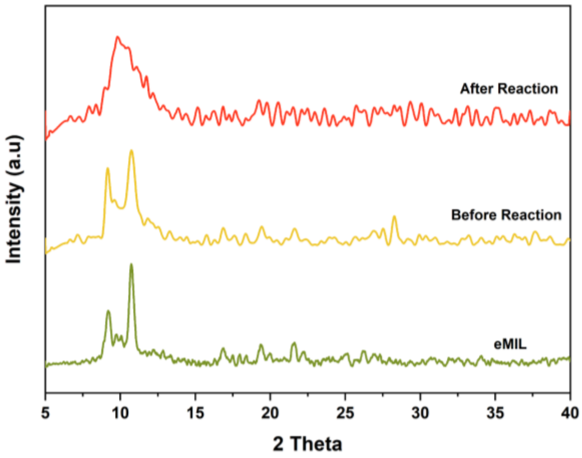


**Figure S9**. PXRD analysis of eMIL before and after pathway infiltration and pathway reaction. Different material quantities were available for the three conditions (eMIL > before reaction > after reaction) leading to worsening signal to noise. Each curve was normalized to its own maximum intensity. Infiltration does not significantly change eMIL crystallinity. After violacein production, crystallinity appears largely conserved but peak broadening may indicate structural changes.

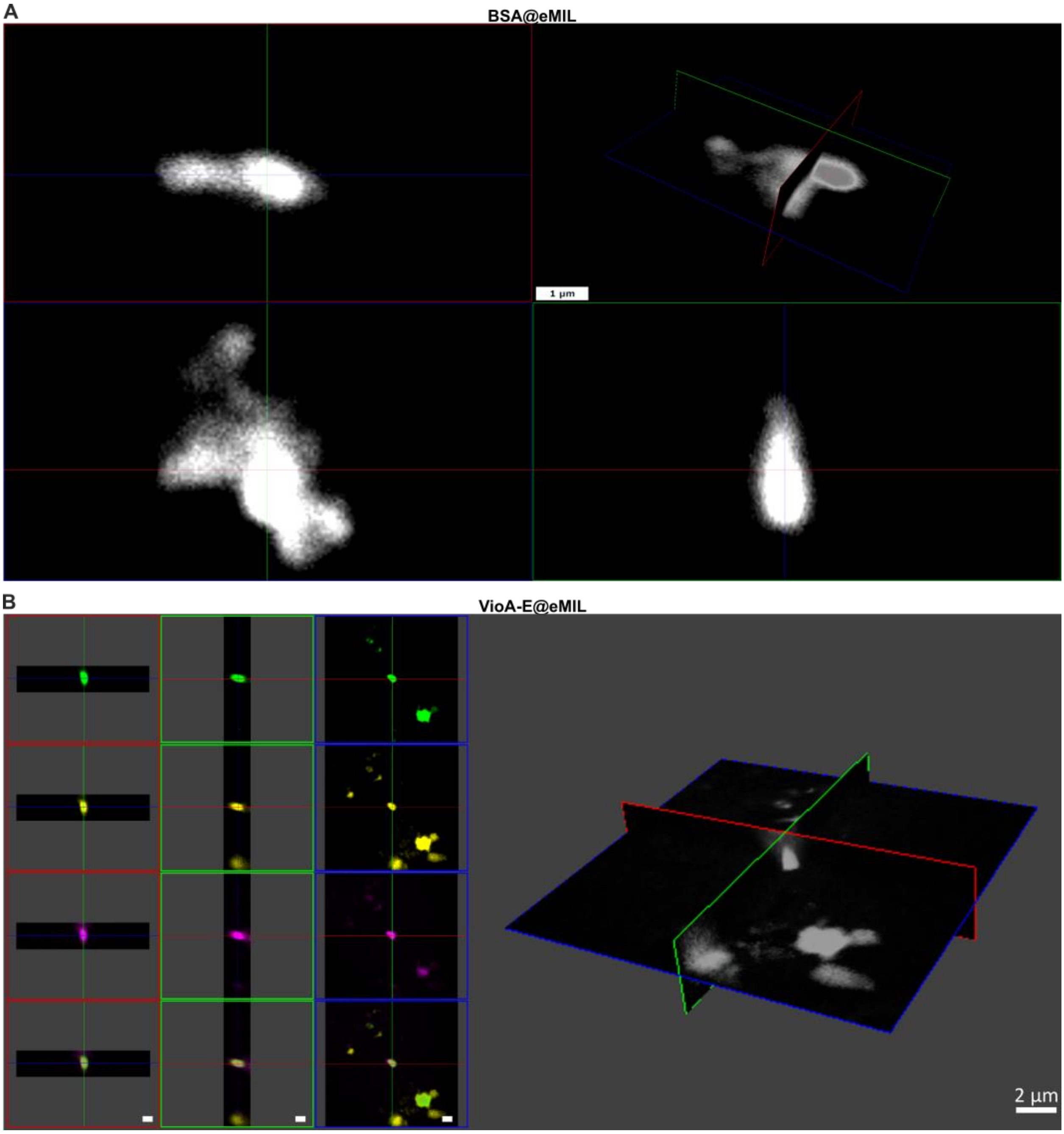


**Figure S10**. Z-stacking confocal fluorescence microscopy of protein-infiltrated eMIL nanoparticles. **A**, FITC-BSA@eMIL. Fluorescently labelled BSA was infiltrated into eMIL at 0.5 mg/ml (BSA) to 2 mg/ml (eMIL) and imaged with 100 x 0.17 µm Z-level steps (see Methods for details). The 3D rendering of a representative cluster comprising several nanoparticles is shown in the upper right with the placement of three orthogonal 3D cross section planes indicated by colors. The remaining images show the three cross sections. In each of these sections, BSA appears homogeneously distributed throughout the complete eMIL volume. Scale bar: 1 µm. **B**, VioA-E@eMIL with different fluorescence labels on three of the five enzymes: AF488-VioB (green), AF594-VioC (yellow), AF647 (VioE) comprising the largest (VioB) and smallest (VioE) enzyme as well as the potentially higher oligomeric VioC. The remaining two enzymes were co-infiltrated without fluorescent label. XYZ acquisitions of each encapsulated protein/dye are shown in separate rows. AF488, AF594, and AF647 were respectively excited at 488, 561, and 633 nm and detected at 500–550 nm, 570–620 nm, and 650–700 nm with HyD detectors in line-sequential mode to prevent bleed-through. A merged 3D grayscale rendering is shown on the right indicating the placement of 3D cross sections. Z-stacks consisted of 50 slices with a 0.14 µm step. Scale bar: 2 µm.

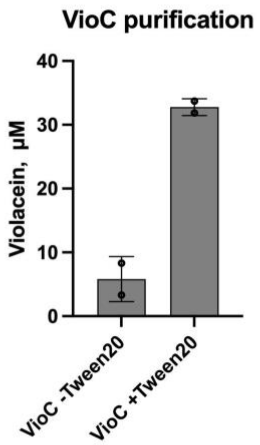


**Figure S11**. Effect of Tween20 on VioC performance. The addition of Tween20 during VioC purification increased the yield of violacein from infiltrated VioA-C@eMIL. Violacein was quantified by UHPLC-MS/MS. Data points of two replicates are shown along with standard deviation.

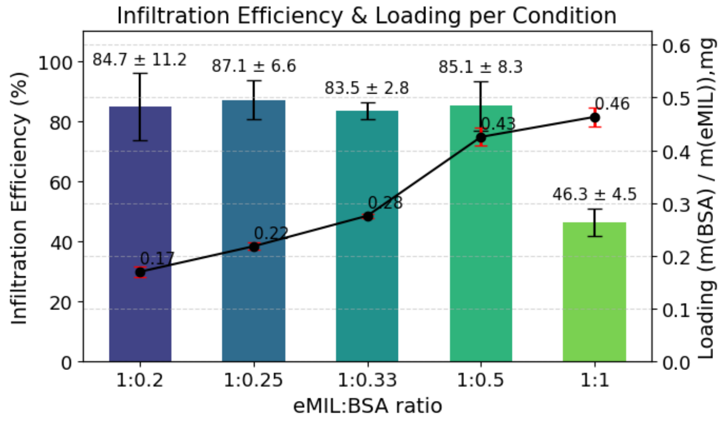


**Figure S12**. eMIL loading capacity and loading efficiency tested with BSA. Varying concentrations of BSA were infiltrated overnight with 2 mg/ml eMIL (glass vials, stirring at 750 rpm, 20 mM Tris, 100 mM KCl, pH=8.0). Supernatant [BSA] was again determined by Bradford assay calibrated against BioRad BSA standards. The standard infiltration condition (1:0.25 eMIL:BSA) is charging the eMIL to less than half of its capacity, confirming similar data recorded for the enzyme mixture in Fig. S4 above.

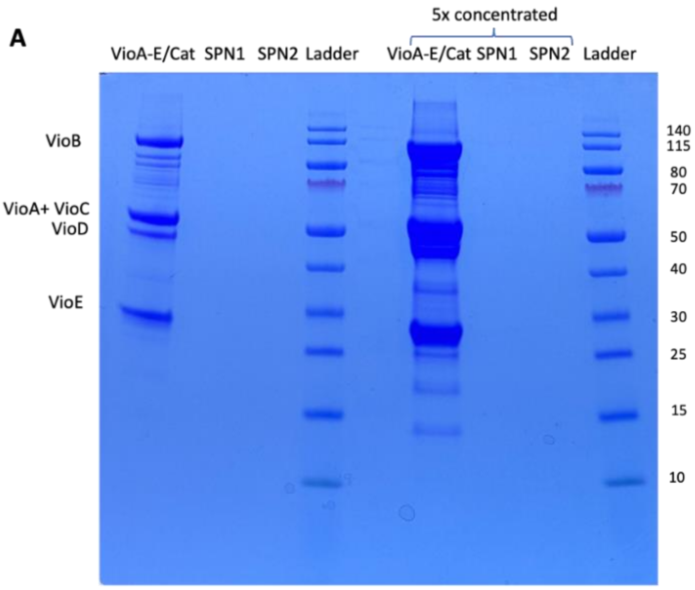


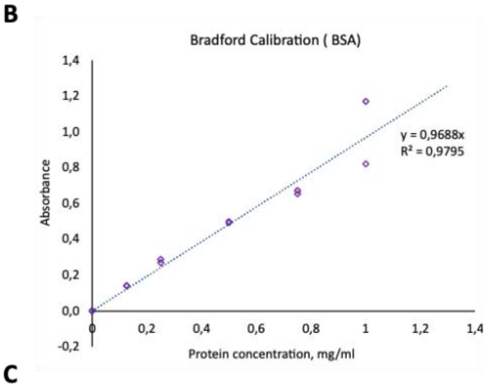


C

| Samples | Concentration in supernatant, mg/ml | Loading efficiency, % |
|---|---|---|
| Supernatant 1 | 0.011±0.008 | |
| Supernatant 2 | 0.007±0.001 | |
| Sum | 0.018±0.009 | 98 |

**Figure S13**. Extended analysis of pathway protein loading into etched MIL. **A**, SDS-PAGE, from left to right: lane 1, original VioA-E/Cat proteins (0.36 mg/ml), lanes 2,3, supernatants from infiltration reactions, 4 ladder, 5 empty (some cross-contamination), 6 - 8 same as 1-3 but after 5-fold concentration of the supernatant with a 10 kDa cutoff spin-concentrator. Absence of any bands indicates full protein infiltration into eMIL. **B**, Bradford assay calibration against BSA. **C**, loading efficiency as determined by Bradford assay on supernatants shown in A.

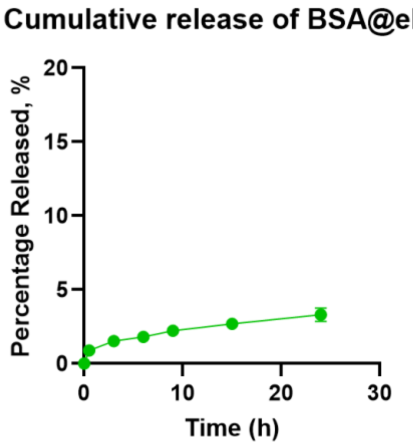


**Figure S14**. In vitro release of FITC-labeled BSA from eMIL. See method section for details. Less than 4% of the infiltrated BSA was released during repeated washes over a 24 h time period.

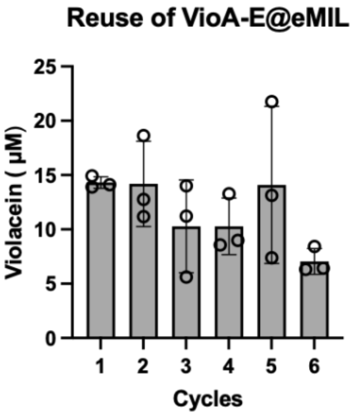


**Figure S15.** Violacein pathway reuse over several cycles of biosynthesis. Violacein recovery from 100 μg of VioA-E@eMIL undergoing 6 cycles of 2h reactions (catalase provided in solution).

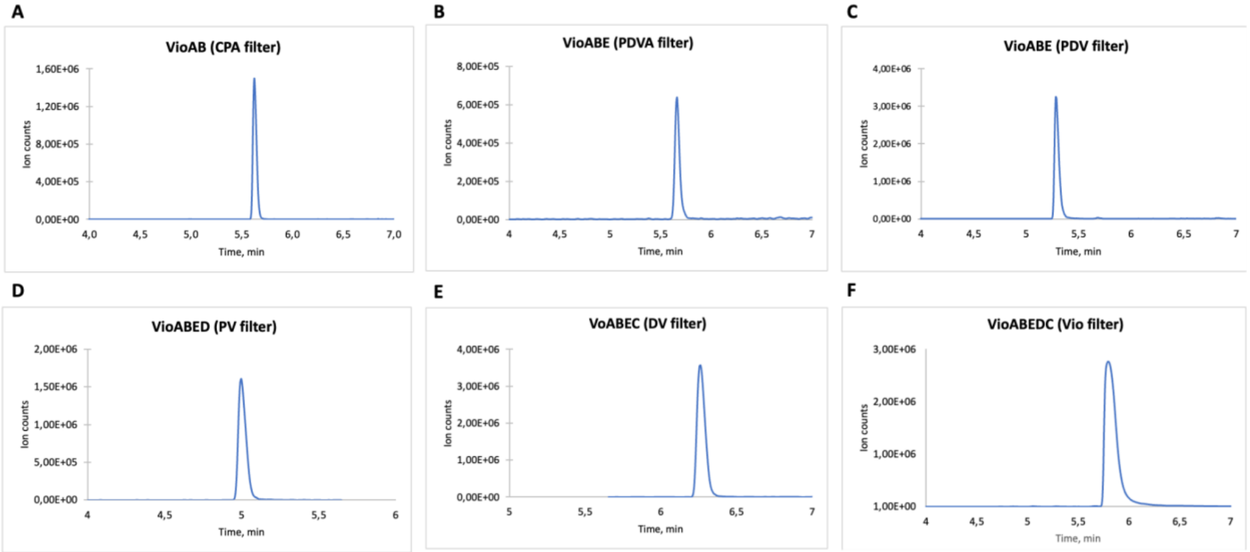


**Figure S16**. Detection of violacein pathway intermediates and side products from incomplete pathways. Shown are UHPLC-MS(/MS) traces after application of SIM or SRM filters. **A**, Chromopyrrolic acid (CPA) produced from VioA+VioB; **B-C**, Prodeoxyviolaceinic acid (PDVA) and prodeoxyviolacein (PDV) produced from VioA+VioB+VioE; **D-E**, Proviolacein (PV) and deoxyviolacein (DV) produced from VioA+B+E+D and VioA+B+E+C, respectively; **F**, Violacein (Vio) produced from full pathway VioA+B+E+D+C. A-E are SIM counts, Vio (F) was detected by SRM (MS/MS), all others by SIM (MS).

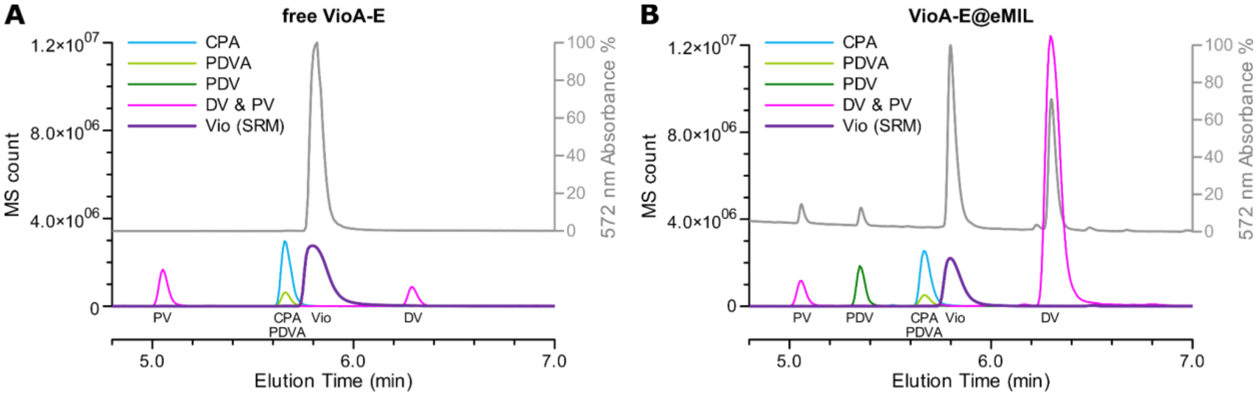


**Figure S17**. UHPLC-MS(/MS) elution profile of VioA-E pathway reactions at the 2h time point before further optimization. **A**, standard reaction (0.36 g/l) of free enzymes in solution. **B**, Infiltrated pathway VioA-E@eMIL with 0.36 g/L total enzyme in 2 g/l eMIL. The 572 nm UV absorbance is shown for comparison. The infiltrated pathway features a generally higher proportion of side products. In particular PDV and DV are increased over the free reaction. However, the UV trace suggests that violacein is still the main product. Please note, as the MS analysis was performed in negative ion mode, higher SIM counts are to be expected for the intrinsically negatively charged CPA and PDVA. More generally, MS counts cannot be readily compared between different compounds or even sets of experiments performed at different times. Elution times may vary with variations in instrument setup (delay volumes).

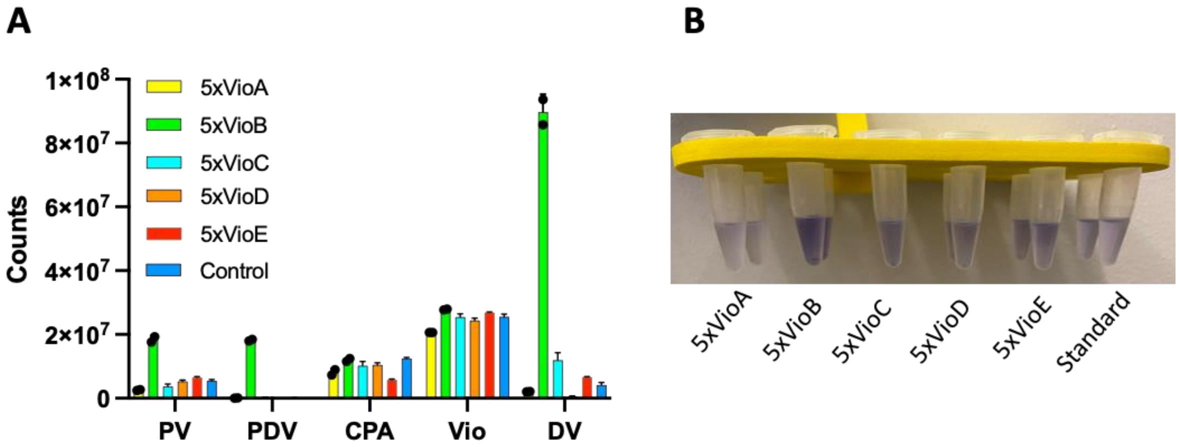


**Figure S18**. The effect of varying enzyme concentration on the performance of the pathway. **A**, Violacein production in solution when individual VioA-E enzymes are increased 5-fold while the five others remain constant. Shown are SRM (MS/MS) counts of violacein and SIM (MS) counts of all other pathway side products. Note that ion counts for a given concentration differ between different molecules precluding direct quantitative comparisons between compounds. Furthermore, detection of Violacein by SRM (MS/MS) rather than regular MS increases specificity but leads to lower counts. **B**, Visual evidence of violacein or deoxyviolacein production for the above reactions (two replicates each). Abbreviations: Chromopyrrolic acid (CPA), prodeoxyviolacein (PDV), proviolacein (PV), deoxyviolacein (DV), violacein (Vio).

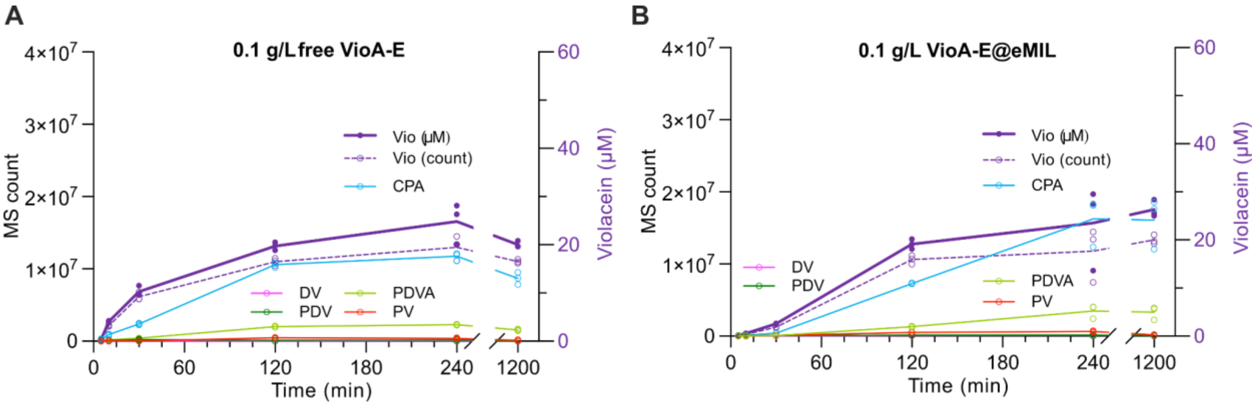


**Figure S19**. Violacein pathway kinetics and side products in five-fold diluted, refined equimolar reactions. Compare with main manuscript Fig. 8 for the full-concentration experiments carried out at the same time. **A**, 0.1 mg/ml total enzyme concentration (0.34 µM each enzyme) in solution. **B**, diluted eMIL reaction with 0.1 mg/ml total enzyme in 0.4 mg/ml eMIL. As before, violacein (purple) concentration was quantified by SRM calibrated against standard (Fig. S20). The SIM count of violacein is given separately (dotted line) for comparison with the SIM signal of the other compounds.

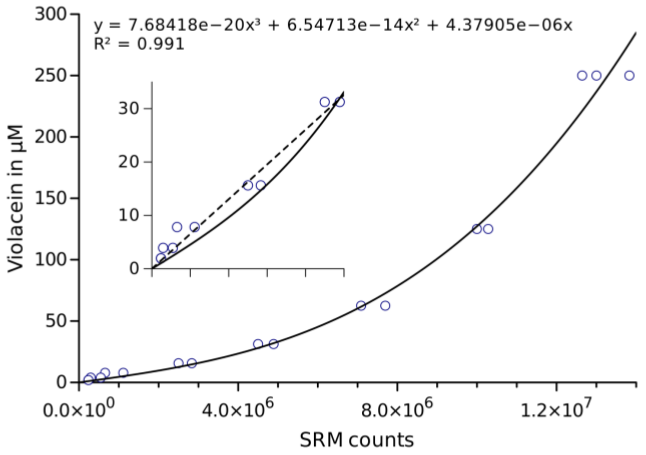


**Figure S20**. Violacein calibration curve for UHPLC-MS/MS experiments. The SRM area under the peak was recorded for violacein standard solutions in a 2:1 methanol:buffer mixture. While the signal was linear for concentrations until 30 µM (inset, broken line), a more complex regression curve (solid line, equation) was used to cover the full concentration range.

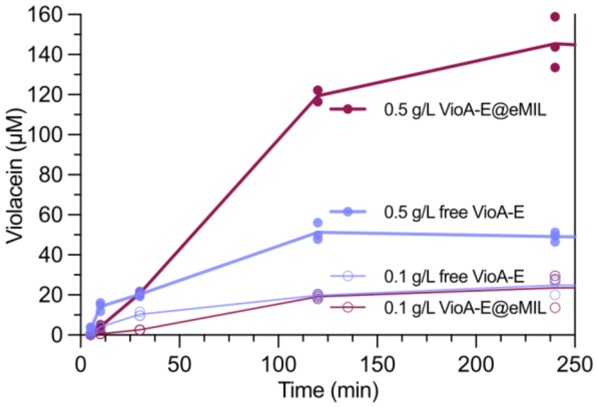


**Figure S21**. Violacein production kinetics from free and infiltrated pathways at different concentrations in direct comparison. This is a re-plotting of the data shown in Fig. 8 and Fig. S19. Pathways in solution (free) show a high initial production rate but decelerate over time. Pathway nanoreactors feature a longer lag phase without initial product burst but then sustain Violacein production for much longer.

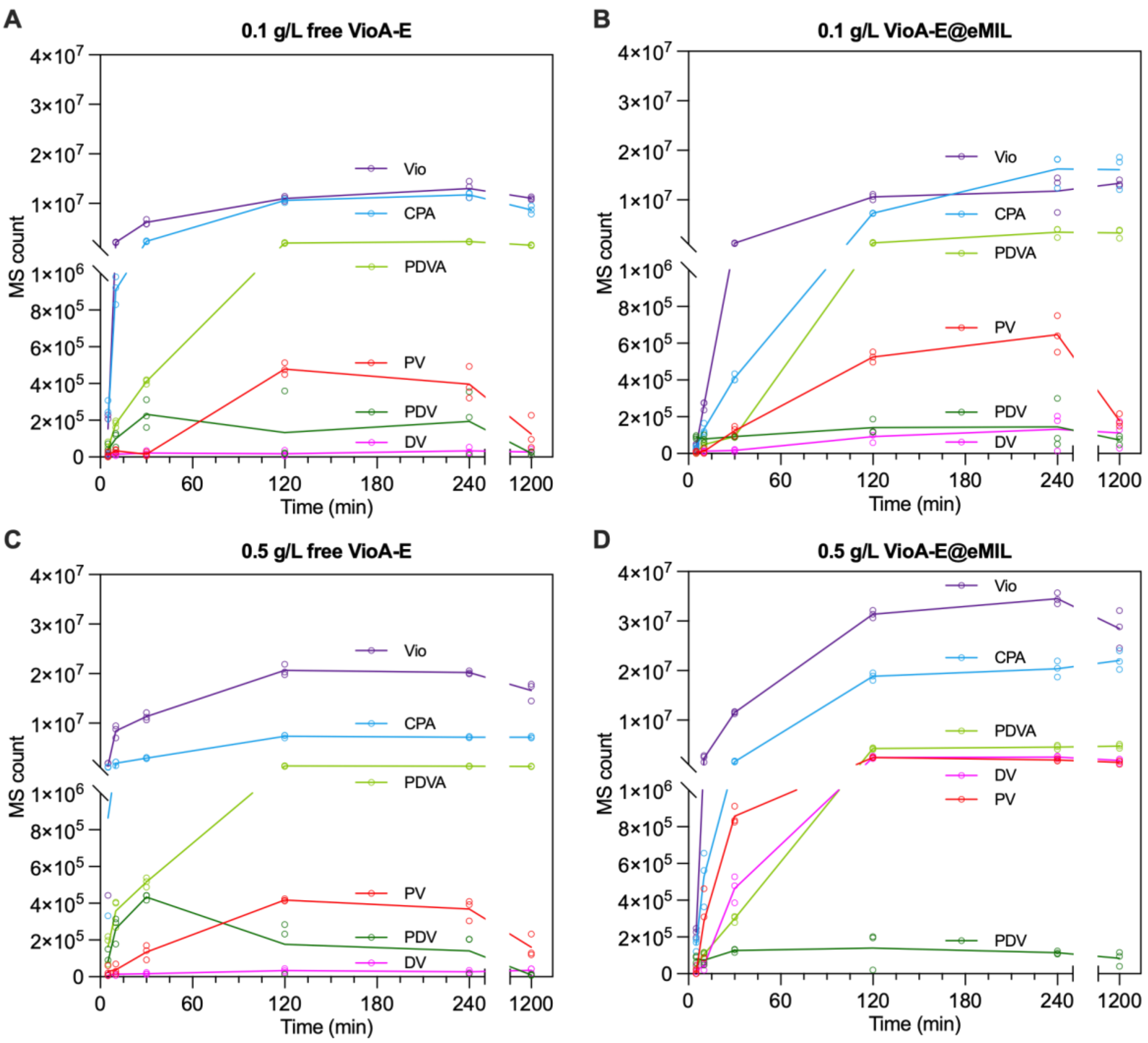


**Figure S22**. Pathway kinetics zooming in at lower abundance side products. This is a re-plotting of Fig. 8C-D in the main manuscript and Fig. S19 above but with the bottom part of the Y-axis enlarged while the top part has been compressed. **A**, 0.1 mg/ml total enzyme concentration (0.34 µM each) in solution. **B**, 0.1 mg/ml total enzyme in 0.4 mg/ml eMIL. **C**, 0.5 mg/ml total enzyme concentration (1.7 µM each) in solution. **D**, 0.5 mg/ml total enzyme in 2 mg/ml eMIL. Note as before that SIM counts cannot be directly translated to absolute concentrations when comparing different compounds. Reduction of PDV and PV concentrations over time may indicate non-enzymatic reaction to (deoxy/oxo/)chromoviridans.

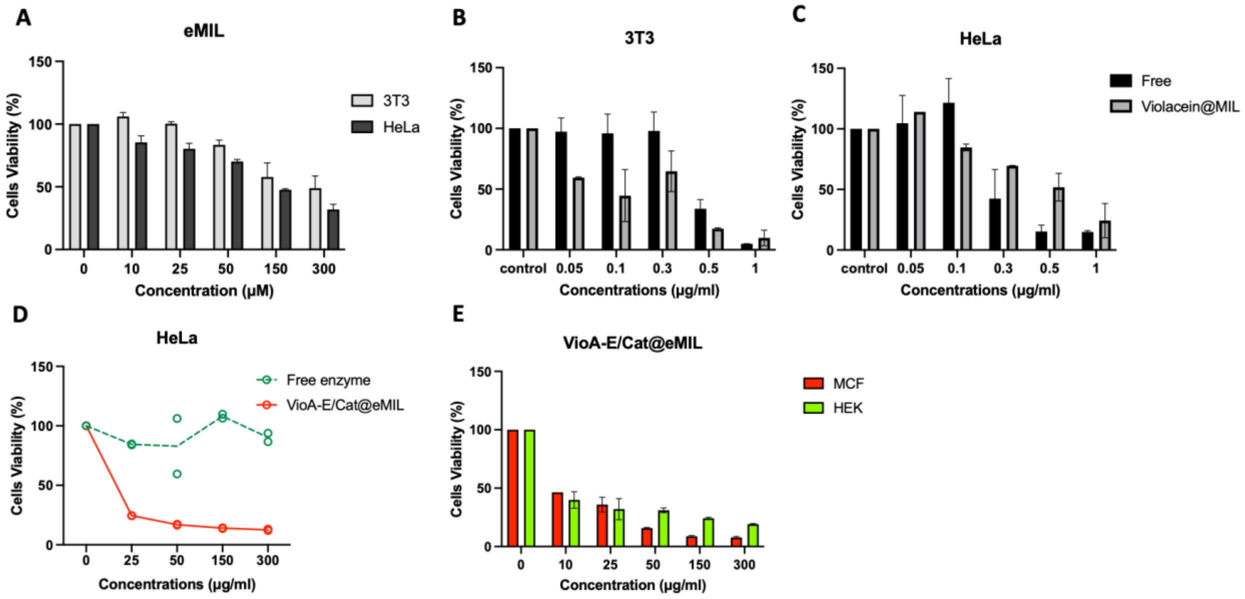


**Figure S23.** Cell viability and internalization studies of eMIL. **A,** Toxicity of eMIL alone in 3T3 and HeLa cell lines. Cell viability of 3T3 (**B**) and Hela cells (**C**) at different concentrations of free violacein (free) and compared to the same concentrations of violacein infiltrated into a constant amount of eMIL (Violacein@MIL). **D,** Cell viability of Hela cells at different concentrations of VioA-E/Cat@eMIL and a control free enzymes **E,** Cell viability of MCF-7 and HEK cells incubated with different concentrations of VioA-E@eMIL, Cell viability was assessed with the CCK assay and normalized to control cells without treatment. Error bars report the standard deviation from three technical replicates.

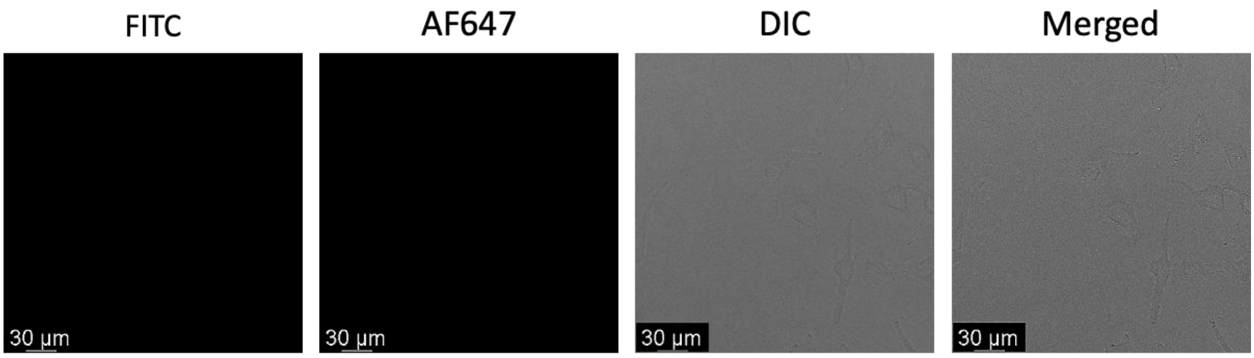


**Figure S24**. Free BSA control for Fig. 6. Confocal fluorescence microscopy of HeLa cells incubated with free BSA-FITC and BSA-AF647 demonstrates no cellular uptake of BSA by itself.

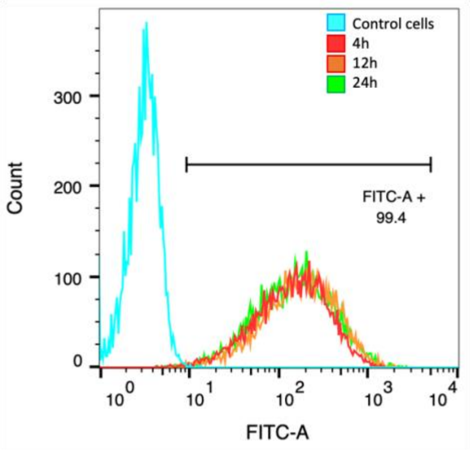


**Figure S25**. Flow cytometry analysis of Hela cells treated with FITC-BSA@eMIL. Hela cells were incubated for 4h, 12h, or 24h with FITC-labeled BSA infiltrated into eMIL. Over 99% cells were FITC-positive already following 4h of incubation, indicating rapid and efficient uptake.

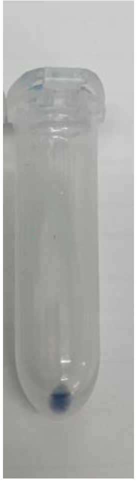

**Figure S26**. HeLa cell pellet after intracellular VioA-E/Cat@eMIL pathway reaction. The violacein product remained tightly associated with the cell pellet, but could be extracted with chloroform : methanol (2:1).
